# Modeling Allelic Diversity of Multi-parent Mapping Populations Affects Detection of Quantitative Trait Loci

**DOI:** 10.1101/2021.07.14.452335

**Authors:** Sarah G. Odell, Asher I. Hudson, Sébastien Praud, Pierre Dubreuil, Marie-Helene Tixier, Jeffrey Ross-Ibarra, Daniel E. Runcie

## Abstract

The search for quantitative trait loci (QTL) that explain complex traits such as yield and flowering time has been ongoing in all crops. Methods such as bi-parental QTL mapping and genome-wide association studies (GWAS) each have their own advantages and limitations. Multi-parent advanced generation intercross (MAGIC) populations contain more recombination events and genetic diversity than bi-parental mapping populations and reduce the confounding effect of population structure that is an issue in association mapping populations. Here we discuss the results of using a MAGIC population of doubled haploid (DH) maize lines created from 16 diverse founders to perform QTL mapping. We compare three models that assume bi-allelic, founder, and ancestral haplotype allelic states for QTL. The three methods have different power to detect QTL for a variety of agronomic traits. Although the founder approach finds the most QTL, there are also QTL unique to each method, suggesting that each model has advantages for traits with different genetic architectures. A closer look at a well-characterized flowering time QTL, qDTA8, which contains *vgt1*, suggests a potential epistatic interaction and highlights the strengths and weaknesses of each method. Overall, our results reinforce the importance of considering different approaches to analyzing genotypic datasets, and show the limitations of binary SNP data for identifying multi-allelic QTL.9

## Introduction

The study of quantitative genetics requires the ability to link differences in phenotype to genotypic variation. Natural and artificial selection act on phenotypes, but only genetic variation will result in changes in population means. Maize presents an excellent model organism to study quantitative genetics due to the combination of extensive genetic and phenotypic resources and the ability to create mapping populations. In addition, maize is one of the most widely produced crops in the world and is a major source of calories for millions of people. Decades of research into maize genetics have resulted in the identification of many quantitative trait loci (QTL) that explain variation in phenotypes such as yield, flowering time, and plant height (Buckler *et al*. 2009; Wang *et al*. 2006; Wallace *et al*. 2014; Beavis *et al*. 1991; Steinhoff *et al*. 2012). Such traits are extremely agronomically important, but are also crucial in terms of fitness and local adaptation.

Researchers have discovered large-effect QTL for a number of agronomic traits in maize through the use of different types of mapping populations (Huang *et al*. 2015). Any choice of mapping population comes with both advantages and limitations. In particular, different types of populations tend to vary in two main characteristics: (1) their ability to capture genetic diversity and (2) their power to detect QTL of small effect. Multi-parent Advanced Generation Intercross (MAGIC) populations have been used in breeding to increase the genetic diversity included in a mapping population compared to biparental populations (Huang *et al*. 2012; Dell’Acqua *et al*. 2015; Highfill *et al*. 2016; Aylor *et al*. 2011; Kover *et al*. 2009; Pascual *et al*. 2015). Compared to genome-wide association panels, MAGIC populations have more power to detect low frequency alleles and can better compare allelic effects between haplotypes because the crossing scheme increases the frequency of all parental alleles to be approximately equal. Simulations of an 8-parent MAGIC population showed that with a sample size of 300 lines, QTL accounting for 12% of variance could be detected with a power of 82% averaged across minor allele frequencies (Dell’Acqua *et al*. 2015). Lastly, a MAGIC population avoids confounding due to population structure that is encountered with genomewide association studies (GWAS) because of the break-up of genome-wide LD.

In this study, we used a MAGIC population of 344 doubled-haploid lines derived from 16 inbred maize parents developed by Biogemma to understand how different quantitative genetic models can impact the identification of QTL. This MAGIC population has an intermediate number of founders compared to the 8-parent (Dell’Acqua *et al*. 2015) and 24-parent (Liu *et al*. 2020) MAGIC populations that have been created. Simulations suggest it should have comparable power to larger nested mapping populations (Dell’Acqua *et al*. 2015; Yu *et al*. 2008). Our population differs from all of these in its use of doubled haploids (DH) instead of recombinant inbred lines. For these reasons, the Biogemma MAGIC population has great potential to reveal new insights into the genetic control of quantitative traits in maize.

In addition to the choice of mapping population, the choice of how to represent genetic information through association and QTL mapping can impact the power of a study to detect and analyze QTL. A bi-allelic model for QTL, often used in GWAS, assumes that a single causal variant at a locus explains the phenotypic variation in the population. Most commonly, genetic variation is represented as bi-allelic SNPs which segregate in the population, with each individual possessing either a reference or alternate allele. With this bi-allelic model, hereafter referred to as *GWAS_SNP_*, each marker’s effect captures the total effect of all variants statistically linked (or correlated) with this marker, generally a small local region of a chromosome.

An alternative model for the allelic state of QTL can be used in multi-parent populations, where we assume that each founder contributes its own allele. In this model, rather than looking at individual SNPs, we test whole chromosomal segments along which the founder state is constant for all individuals in a population. As a result, QTL are multi-allelic, with the number of alleles equal to how many founders were used in the making of the population. We will refer to this founder model hereafter as *QTL_F_*.

The assumption that each founder contains a functionally and evolutionarily distinct haplotype in a genomic window, although likely for biparental mapping populations, becomes increasingly unlikely as the number of founders increases. This is because the founders of a population likely share ancestral haplotypes through identity-by-descent (IBD). A third allelic model takes into account shared ancestral haplotypes between founders. This model, hereafter referred to as *QTL_H_*, allows the number of alleles at each site to vary anywhere from one to the total number of founders (here 16), based off of the number of ancestral haplotypes at that site. This has the potential to increase statistical power compared to the *QTL_F_* model by reducing the number of parameters.

Here we present a maize MAGIC population derived from 16 parents and discuss the performance of three different models for representing allelic states: bi-allelic, founder, and ancestral haplotype allelic models for detecting QTL. Using *vgt1*, a well-characterized flowering time QTL with a strong candidate causal variant that is variable in the population, we demonstrate differences between the three methods and explore potential interactions between *vgt1* and other genetic variation in the population.

## Materials and Methods

### Mapping Population

The MAGIC population was derived from 16 inbred maize parents representing the diversity of European flint and U.S. dent heterotic groups. The 16 founder lines were crossed in a funnel crossing scheme, and then the resulting synthetic population was intercrossed for 3 generations with around 1600 individuals per cycle (Figure 1A) (Supplemental File 1). Finally, 800 lines were selected from the synthetic population to create doubled haploids (DH), resulting in 550 MAGIC DH lines at the end of the process. The MAGIC DH lines were crossed to a tester MBS847 to produce 344 hybrids (Figure 1A). Due to variation in flowering time, a subset of the lines could not be crossed to the tester (Supplemental File 1).

**Figure 1.**
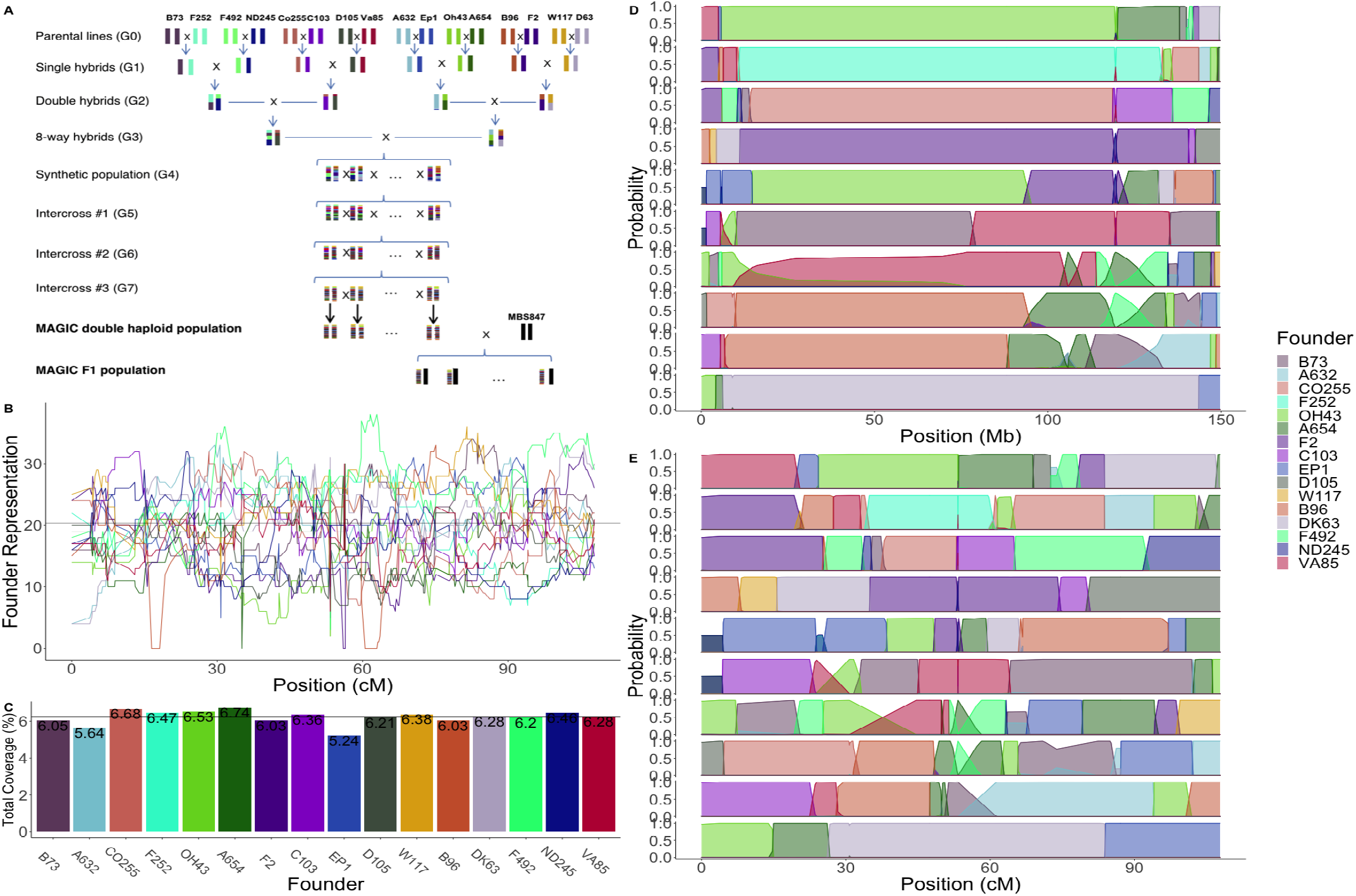
Structure, diversity, and founder representation of the MAGIC population. (**A**) The crossing scheme of the MAGIC population. (**B**) Coverage of each founder across the population on Chromosome 10. The horizontal black line represents the expected number of lines per founder with equal distribution (21.5). (**C**) Total coverage of each founder across the population as a percentage. The black line shows the expectation of equal distribution (6.25%) (**D**) Founder probabilities for 10 MAGIC DH lines on chromosome 10 in physical distance. (**E**) Founder probabilities for 10 MAGIC DH lines on chromosome 10 in genetic distance.

### Genotyping

The 16 founder lines and the MAGIC DH lines were all genotyped with the Affymetrix 600K Axiom SNP array (Unterseer *et al*. 2014), resulting in genotype data for 517,769 SNPs. A total of 503,902 SNPs were used after filtering out invariant sites and sites that were not located on autosomal chromosomes according to the B73 AGPv4 reference genome (Jiao *et al*. 2017).

### Phenotype Data

The MAGIC F1 plants were phenotyped in five different field locations in four different years, resulting in six distinct environment-years (Hudson *et al*. in prep). The environments represent a range of latitudes and water stress, from vegetative and flowering water deficit (Nerac 2016) to optimum well-watered conditions (Graneros 2015). In each environment we grew a minimum of 292 and a maximum of 309 of the DH lines. Each genotype was grown with two replicates in each environment. In all environments, seven traits were measured: grain yield (GY), plant height (PH), female flowering date (DTS), male flowering date (DTA), thousand kernel weight (TKW), and harvest grain moisture (HGM), and anthesis-silking interval (ASI) (Hudson *et al*. in prep). For each of the lines we calculated best linear unbiased predictor (BLUP) scores for all seven phenotypes, combining measurements from all environments to get estimates of the genetic contribution to the phenotype for each MAGIC line (Aulchenko *et al*. 2007).

### Calculation and Validation of Founder Probabilities

We used the package R/qtl2 (Broman et al. 2019) to determine founder probabilities of the MAGIC DH lines using the 600K genotype data and the cross type “riself16”. SNPs used as markers for the *QTL_F_* approach were filtered based on linkage disequilibrium using an iterative approach where a SNP was dropped if the *R*^2^ value of probabilities between it and the previous SNP was greater than 0.95. After filtering, a total of 4,578 sites were kept.

Due to the fact that the actual crossing scheme and the cross type input into R/qtl2 differed (DH lines rather than RILs), we wanted to assess the accuracy of the founder probabilities. This was done by simulating lines using the actual crossing scheme and assessing the performance of the calc_genoprobs function of R/qtl2 in correctly identifying the founder genotype (Figure 1D & E). We developed an R package (R Core Team 2017), *magicsim* (https://github.com/sarahodell/magicsim) to simulate the lines using the maize consensus genetic map from (Ogut *et al*. 2015) to generate approximate recombination rates across the chromosome. We simulated 100 MAGIC populations constituting 344 lines and assessed founder assignment accuracy as the average percentage of SNPs where the predicted founder was the same as the actual founder.

### Test for Equal Representation of Alleles

The number of lines that had a probability of a particular founder greater than 0.8 were used as an approximation of the number of lines that had that founder at a site. This observed count was compared to a null expectation of 1/16 for equal distribution across lines (approximately 21 lines per founder)(Figure 1B). We performed a *χ*^2^ test for each site to determine if founder counts significantly deviated from null expectation. We obtained a 5% significance threshold using the *χ*^2^ obtained from founder counts in 100 simulated populations. The *χ*^2^ tests from simulated lines were done using reconstructed genotype data pulled from the 600K genotype data of the 16 founders. We then used the same methods of calculating founder probabilities with R/qtl2 used with the actual population. Due to the fact that we used the inferred founder identities of the simulated lines, rather than the known founder identities, the null distribution of p-values generated from *χ*^2^ tests of the simulated populations incorporated uncertainty of founder assignment.

### Calculation of Identity-by-Descent and Haplotype Probabilities

The identification of regions of shared genetic sequence between founder pairs allows collapsing of founders into ancestral haplotypes. IBD was measured from the 600K SNP data of the founders using the software RefinedIBD (Browning and Browning 2013) with a sliding window of 10 cM and a minimum IBD segment length of 0.2 cM. The resulting segments of pairwise IBD between each of the 16 founders were used to identify distinct haplotype blocks. We did this by moving along the chromosome, starting a new haplotype block when a segment of pairwise IBD between founders started or ended (Figure S1). Then, within blocks, we grouped all founders that were in IBD with one another into a haplotype and summed the founder probabilities to obtain haplotype probabilities.

In certain instances, the pairs of founders that were in IBD with one another in a particular haplotype block formed an incomplete graph, where not all founders were in IBD with all other founders (Figure S1). For example, from the results of RefinedIBD, an incomplete haplotype graph of three founders would have founder A in pairwise IBD with both founder B and founder C, but founder B and C not in pairwise IBD. For the sake of simplicity, we assumed that all founders in a haplotype were in IBD with one another (we called B and C as in IBD). However, it is important to note that haplotypes called here may still possess genetic differences between founders, with some founders being more different than others.

Markers for the *QTL_H_* mapping approach were filtered for LD using an iterative approach similar to *QTL_F_*: for all haplotype blocks with the same number of distinct haplotypes, a SNP was dropped if the correlation of probabilities between it and the previous included SNP was greater than 0.95. After filtering, a total of 11,105 sites were kept to represent haplotype blocks in the MAGIC DH lines.

### Association and QTL Mapping

The R package GridLMM (Runcie and Crawford 2019) was used to run association mapping using the three different methods of representing the genotype data. The function GridLMM_ML was used with the “ML” option. The following three models were approximated by fitting each locus independently. The three methods differed in the *X* matrix used in the mixed linear model.

The bi-allelic model (*GWAS_SNP_*) was

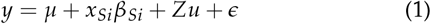

where *y* is the response variable, *μ* is the global mean, *x_Si_* is an *n* x 1 genotype vector for SNP *i* with reference and alternate alleles represented as 0 and 1, respectively, *β_Si_* is the effect size of the alternate allele, *Z* is the design matrix, 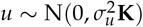 is the random effects of markers across the rest of the genome using the genomic relationship matrix, *K*, and *ϵ* is the error.

The founder model (*QTL_F_*):

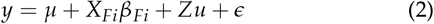

where *y* is the response variable, *μ* is the global mean, *X_Fi_* is a *n* x *f* − 1 matrix for marker *i* and *X_fni_* is the probability that at site *i*, individual *n* was derived from founder *f*, *β_F_i*, is the effect size of each founder allele, *Z* is the design matrix, 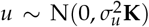 is the random effects of markers across the rest of the genome using the genomic relationship matrix, *K*, and *ϵ* is the error.

The ancestral haplotype model (*QTL_H_*):

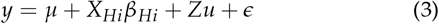

where *y* is the response variable, *μ* is the global mean, *X_Hi_* is an *n* x *h* − 1 matrix for marker *i* and *x_hni_* is the probability that at site *i*, individual *n* has haplotype *h*, *β_Hi_*, is the effect size of each haplotype allele, *Z* is the design matrix, 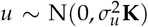 is the random effects of markers across the rest of the genome using the genomic relationship matrix, *K*, and *ϵ* is the error.

Significance cutoffs for p-values were obtained using permutation testing, taking the 5% cutoff from 1000 permutations where genotypes were randomized relative to phenotypes for each method.

### Model Comparison

The results of the three models were compared using two main criteria: (i) presence or absence of identified QTL peaks and (ii) the size of QTL support intervals. QTL support intervals were determined by identifying the most significant SNP for a QTL peak and demarcating the left and right bounds of the QTL as the left-most and right-most SNPs within a 100Mb window centered on the highest SNP that have a -log10(p-value) that is 2 log10(p-values) below that of the highest SNP. The detection of QTL was compared across the three methods for each phenotype. A QTL was said to be identified across models if the QTL support interval for that QTL overlapped. The effect of the model used on the size of QTL support intervals was investigated using the QTL which were identified by all three methods (n=26).

The support interval size response variable was represented both in terms of physical distance (Mb) and genetic distance (cM).

### Estimation of Effect Sizes

We used the R package *lme4qtl* to calculate standard errors of effect sizes relative to the population mean (Ziyatdinov *et al*. 2018). For the *QTL_F_* model, effect sizes were dropped for individual founders at some sites if there where fewer than 5 MAGIC lines that had a probability greater than 0.8 for one of the founders. This same filtering was done with *QTL_H_* effect sizes for sites with low representation of particular haplotypes. This was to ensure that effect sizes for individual founders and haplotypes could be effectively estimated. We confirmed that effect sizes calculated by GridLMM and lme4qtl matched one another, with the correlations in effect sizes between the two methods greater than 0.99.

### Tests for Epistasis

We ran a genome scan for epistatic interactions with *vgt1*. The probability of the MAGIC lines having the MITE insertion at *vgt1* was calculated by summing the founder probabilities for all founders that have the *MITE^+^* allele at the site closest to the location of the MITE underlying *vgt1* in the B73 APGv4 assembly found on MaizeGDB (Portwood *et al*. 2018). Lines that had uncertain allelic states at the MITE (0.05 > Pr(MITE^+^) < 0.95) were dropped for the test. Applying a Bonferroni significance threshold adjusted for the number of tests, we tested for epistasis using the 600K genotype data.

We also performed QTL mapping with *QTL_F_* using only the *MITE^+^* MAGIC lines. This was to see if there were any other loci whose effect was only observed in the presence of the MITE. A normal epistatic model could not be fit with founder alleles because there were not enough degrees of freedom to compare each founder. We used the model from Equation 2 using DTA BLUP scores and the 5% significance threshold for DTA.

### Flowering Time Enrichment Test

We used a list of flowering time (FT) genes assembled by (Wang *et al*. 2017) to test for enrichment of FT genes in founder *χ*^2^ peaks. Of the 907 genes, we used 887 which were aligned to chromosomes 1 through 10 in the B73 AGPv4 assembly (Jiao *et al*. 2017). To determine a null distribution, we randomly sampled 887 non-FT genes and counted the number that overlapped with regions within *χ*^2^ peaks. We compared this number to the actual number of FT genes that overlap with *χ*^2^ peaks.

### Data availability

Genotypic data will be made available through FigShare. Phenotypic and environmental data will be made available at FigShare associated with our companion paper, (Hudson *et al*. in prep)(tracking number G3-2021-402688) (https://figshare.com/s/5ee8337defdef63b04ce). Supplemental files are available at FigShare. File S1 contains a detailed description of the crossing scheme used to develop the MAGIC population. File S2 shows Manhattan plots similar to 3 for individual environments. File S3 shows founder effect size plots for *vgt1* similar to 4 for individual environments. File S4 is contains tables of all QTL identified in the study, their locations, and their effect sizes estimates from the three models. Code used to run analyses and to generate simulated data can be found at https://github.com/sarahodell.

## Results

### MAGIC Population

We developed a 16-parent MAGIC population using temperate inbred maize lines representative of the diversity of the Flint and Dent heterotic groups of North America and Europe (Figure 1A). We genotyped 334 MAGIC DH lines from the population with a 600K SNP genotyping array and measured 7 phenotypes across 6 environments from hybrids resulting from crossing the DH lines to an inbred tester, MBS847. Using the phenotype data from the six environments, we calculated BLUP scores for each of the MAGIC lines (Hudson *et al*. in prep). PCA Analysis of the MAGIC lines and the 16 founders and tester showed that the MAGIC lines maintained an expected amount of genetic variation (Figure S2). In addition, the minor allele frequencies of SNPs in the 16 founder compared to in the MAGIC lines suggested that lower frequency SNPs in the founders were brought up in frequency in the MAGIC lines, which aids in the estimation of SNP effect sizes (Figure S3).

### Simulation and Validation of Founder Probabilities

We partitioned the genomes of individual MAGIC lines into segments of ancestry from the 16 founders. This allowed us to determine the predicted contribution of each founder to the population (Figure 1D & E). The founder probabilities determined using R/qtl2 were able to assign founders to the actual MAGIC DH lines with high confidence (>0.80) for 96.7 % of the 10 chromosomes of maize. The median size of recombination blocks was 4.325 Mb and the mean size was 15.761 Mb with a standard deviation of 29.524 Mb. The average number of crossover events per line was 123.7 with a standard deviation of 20.73. Our simulations suggest a very high (*μ* = 99.8%, *σ* = 0.011) assignment accuracy (see Methods). This reinforced our confidence in the founder probabilities obtained from the actual data.

### Identity-By-Descent and MAGIC Haplotypes

In some cases, the model which inferred founder identity in the MAGIC lines had high uncertainty, with probabilities split approximately equally between two founders. We hypothesized that this uncertainty was due to the two founders having very similar genetic sequence at those regions, such that the model struggled to differentiate the two. To assess the genetic similarity of the founders, we calculated pairwise Identity-By-Descent (IBD) between all founders using the software Refined-IBD (Browning and Browning 2013). As expected, areas of uncertainty in founder probabilities of the DH lines were associated with regions of IBD between two or more founder lines in that region of the chromosome (Figure S4).

The results showed that a total of 1.81 Gb (86%) and 1367.5 cM (92.7%) of the genome were in IBD between at least two different founders. The average size of an IBD segment between two founders was 140kb (0.51 cM) with a median of 122kb (0.46 cM). Pairwise IBD segments sizes ranged from 8 kb (0.3 cM) to 673 kb (1.61 cM). For founder pairs that were in IBD with one another, the total percentage of IBD between founders ranged from 0.0018% (F492 and VA85) to 4.39% (B73 and A632), with an average of 0.061%. There were no IBD segments found for 18 of 120 possible pairwise founder combinations. The amount of IBD segments between the 16 founders and the tester, MBS847, was mostly low (ranging from 0.14% of the genome for F2 to 3.7% for B73), with the notable exception of DK63, which was in IBD with MBS847 for 36.5% of the genome. The Neighbor-Joining Tree of the relatedness of the 16 founders and MBS847 recapitulated the IBD results (Figure S5). For a particular founder pair, B73 and A632, there were large segments where the lines shared haplotypes, and the tree placed them very close together. This is consistent with the pedigree of the lines, where A632 was derived from B14, a line from the same heterotic group as B73 (Lorenz and Hoegemeyer 2013).

Due to the widespread Pairwise IBD between the founders, it appeared that many founders shared ancestral haplotypes. Within individual blocks of ancestry, we collapsed founder alleles that were identical by descent into a single haplotype. The genome was broken up into a total of 6,929 haplotype blocks. Of those blocks, approximately 16% of them (1,152) contained at least one haplotype whose pairwise IBD between parents was incomplete, meaning that there was some genetic variation between founders within those haplotypes that was not captured by the haplotype designation (see Methods). The number of unique haplotypes within haplotype blocks varied across chromosomes, ranging from 6 at the lowest to 16 at the highest (Figure 2A & B). The average number of unique haplotypes per haplotype block was 13 (*μ* = 12.85, *σ* = 1.71) (Figure 2B). There was a wide range of haplotype block sizes, with the average physical size of haplotype blocks being 303.7 kb (*σ* = 1.71Mb) (Figure 2C). The largest haplotype block was 39.3 Mb long on chromosome 7, which had 16 unique haplotypes. In genetic distance, haplotype block sizes range from 0 to 3.4 cM, with an average of 0.20 cM and a median of 0.11 cM (Figure 2D).

**Figure 2.**
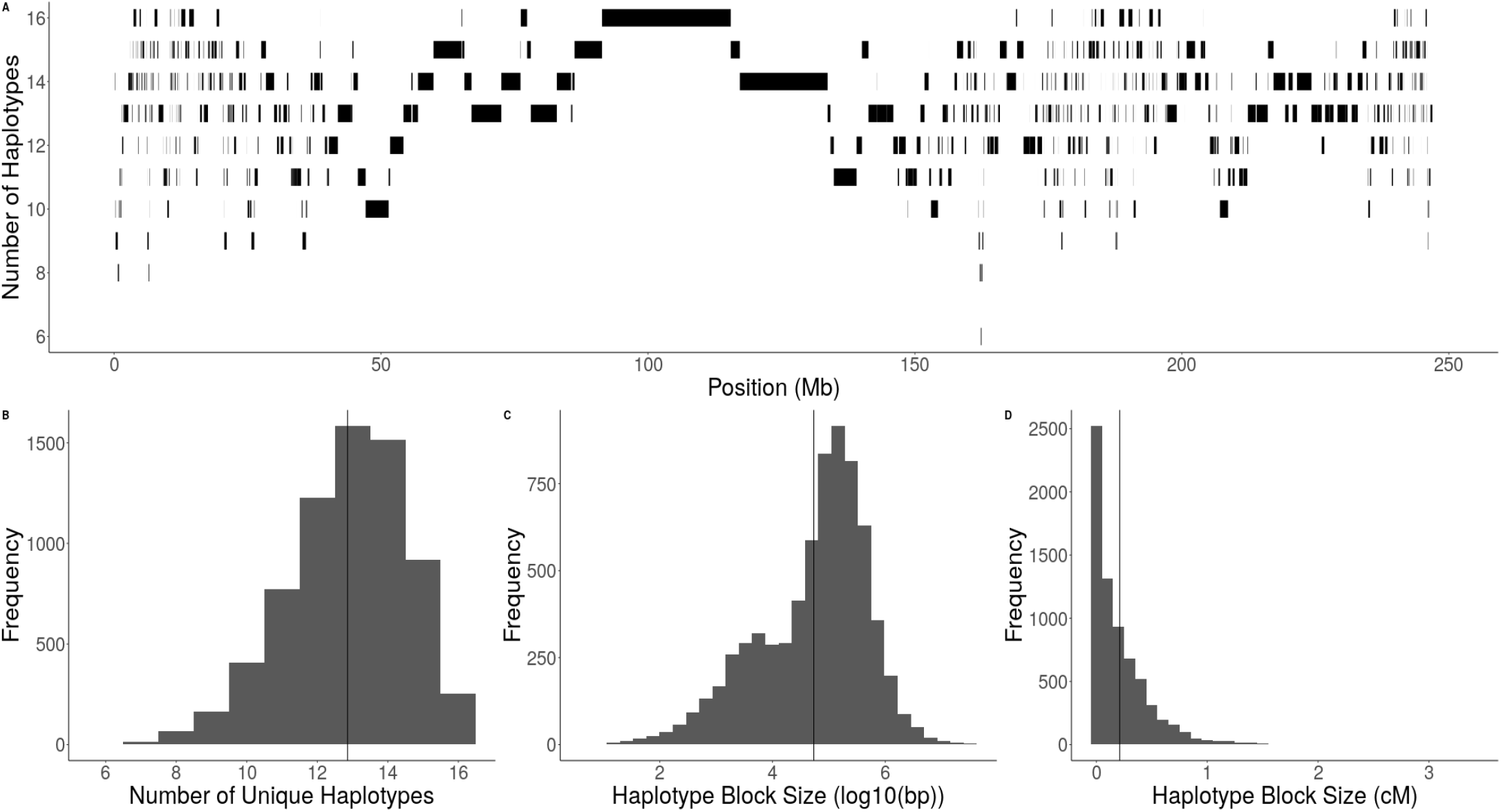
Diversity and Size of Haplotype Blocks. The number of unique haplotypes among the 16 MAGIC founders across the 10 chromosomes of maize (**A**) The number of unique haplotypes per haplotype block and size of haplotype blocks along chromosome 4 in physical distance. (**B**) Distribution of unique haplotypes per haplotype block across the genome. (**C**) Distribution of haplotype block size in physical distance, represented as log10(bp). The black bar represents the average size of 303.7 kb. (**D**) Distribution of haplotype block size in genetic distance, represented as cM. The black bar represents the average size of 0.2 cM.

### QTL Mapping and Association Mapping

We performed association mapping using three models of the allelic state of QTL. The first method, *GWAS_SNP_*, used SNP genotypes obtained from the 600K array, assuming that QTL are bi-allelic. The second method, *QTL_F_*, used probabilities of founder identity in chromosome segments, assuming that a QTL had as many alleles as founders. The third method, *QTL_H_*, used probabilities of haplotype identity in chromosome segments, assuming that a QTL had as many alleles as ancestral haplotypes. We performed QTL mapping separately in each of the 36 environment:phenotype combinations, plus the across-environment averages of the 6 traits, for a total of 42 separate analyses. The three methods varied in their ability to identify QTL. The majority (26 or 57%) of QTL were identified by all three models, with 6 QTL found in both *QTL_F_* and *QTL_H_* and 1 QTL was found in both *GWAS_SNP_* and *QTL_F_*. In addition, each model found unique QTL, with 7, 3, and 3 QTL found in only *GWAS_SNP_*, *QTL_F_*, and *QTL_H_*, respectively (Figure S6).

Next we merged QTL of the same phenotype from different environments based on overlapping support intervals. There were 20 unique across-environment QTL identified from all methods, and 10 of these were found in more than one environment. We found 12 across-environment QTL using BLUPs and 2 of these were not identified in any individual environment. Of these across-environment QTL, 6 (0 unique) QTL were found in Blois, 2014, 7 (1 unique) in Blois, 2017, 7 (3 unique) in Graneros, 2015, 5 (1 unique) in Nerac, 2016, 6 (3 unique) in St.Paul, 2017, and 3 (0 unique) in Szeged, 2017.

Figure 3 shows the Manhattan plots from the three methods for BLUP DTA overlayed with the support intervals of significant QTL for all BLUP phenotypes. Analysis with individual environments identified fewer QTL, but displayed similar patterns (Supplemental File 2). For DTA, all three methods easily identify two large QTLs, *qDTA3-2* on chromosome 3 and *qDTA8* on chromosome 8. These QTL correspond to three previously identified QTL, *vgt3* for *qDTA3-2* and *vgt1* and *vgt2* for *qDTA8*. In addition, there are multiple QTL that are found by only a subset of the models. Using BLUPs, the *GWAS_SNP_* method was able to identify one QTL on chromosome 5 for thousand-kernel weight, *qTKW-5* that was not found in either *QTL_F_* or *QTL_H_*. A second QTL on chromosome 3 for DTA, *qDTA3-1*, was only found with the *QTL_H_* method. A DTA QTL on chromosome 9, *qDTA9* and a harvest grain moisture QTL on chromosome 3, *qHGM3-1* were only found using *QTL_F_*. However, a QTL for DTS with overlapping support intervals to *qDTA9* was found in both *QTL_F_* and *QTL_H_*.

**Figure 3.**
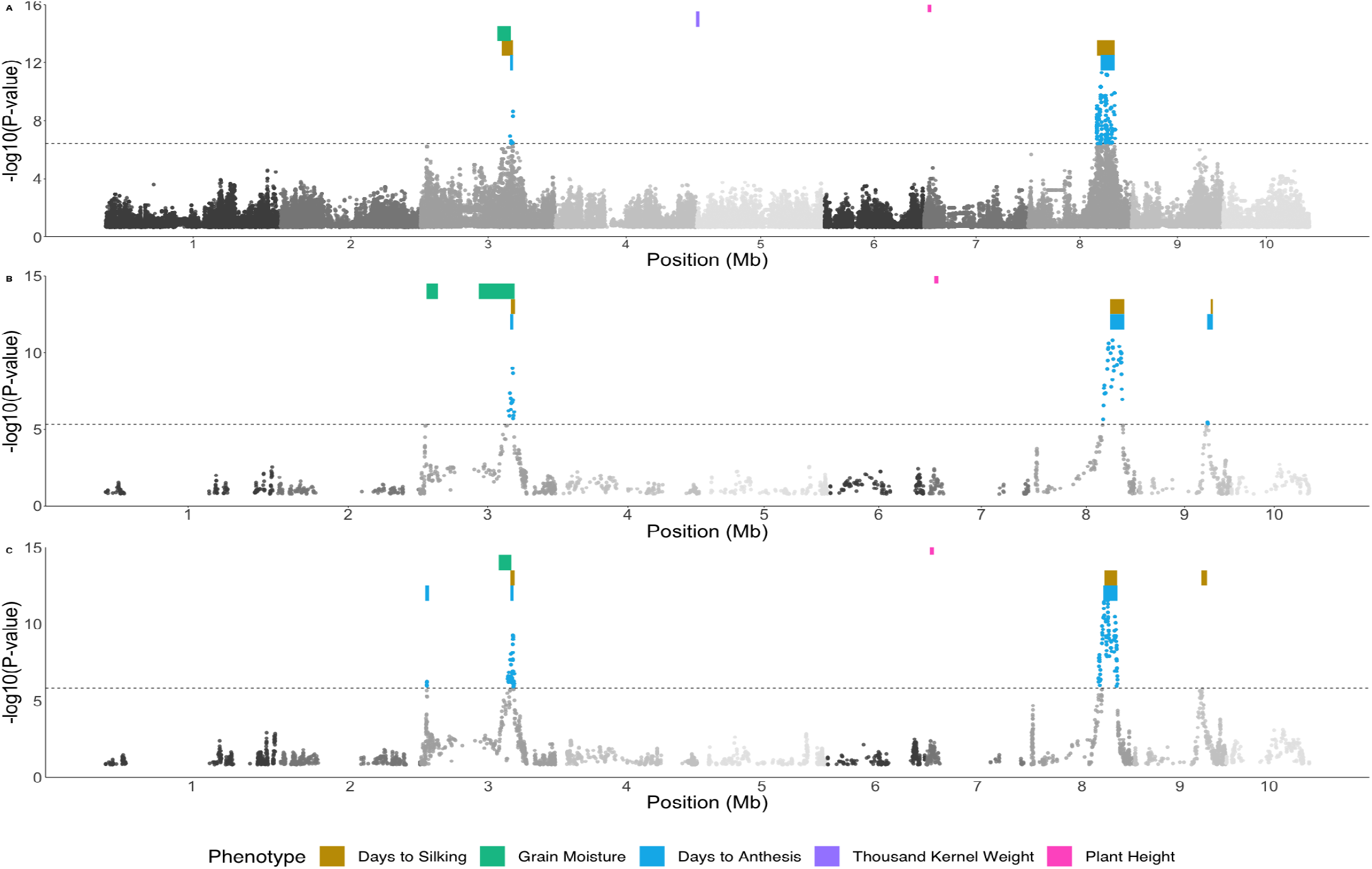
Results of three methods of QTL identification. GWAS results using **(A)** *GWAS_SNP_* **(B)** *QTL_F_*, and **(C)** *QTL_H_*. Colored points represent significant QTL for Days to anthesis (DTA) BLUPs above the 5% threshold from 1000 random permutations, located between -log10(p-value)=5 and -log10(p-value)=6 (dashed line). Colored bars represent QTL support intervals for all other BLUP phenotypes for which a QTL was found, DTS (yellow), HGM (green), DTA (blue), TKW (purple), and PH (pink)

Despite differences in the models, the power to identify and refine the location of QTL was similar across the three methods. *QTL_F_* was able to identify the most QTL, regardless of changes in the significance threshold (Figure S7). Individual QTL that were found in one method at the 5% significance threshold usually became significant in other methods when the 10% threshold was used, indicating that the differences in the ability to detect these QTL between methods is mostly due to differences in power (Figure S7). Nonetheless, there were multiple QTL that were identified in only one method. There were 10 QTL that appeared in only one method, mostly related to grain yield and thousand kernel weight traits.

Support intervals for QTL were determined using a twofold change in the log_10_ p-value of the most significant SNP (see Methods). The size of *GWAS_SNP_* support intervals were significantly larger than *QTL_H_* support intervals in genetic distance (1.55 cM, SE = 0.56, t-ratio = 2.77, p-value = 0.021), but the difference in physical distance was not significant (t-ratio = 1.97, p-value = 0.13)(Table S1). There was no significant difference in the size of QTL support intervals between between *QTL_F_* and *QTL_H_* in either physical and genetic distance. Although on average, the physical and genetic size of *GWAS_SNP_* support intervals were larger than those of *QTL_F_* support intervals, the difference was not significant, perhaps because of a single outlier QTL in *QTL_F_* with a very large support interval (Figures S8 & S9). When the outlier was dropped from the model, the difference between *GWAS_SNP_* and both *QTL_F_* and *QTL_H_* support intervals were significant in both genetic (t-ratio = 2.87, p-value = 0.017; t-ratio = 3.17, p-value = 7.4e-3) and physical distance (t-ratio = 2.91, p-value = 0.015; t-ratio = 2.68, p-value = 0.027).

### *Variation around* vgt1

One notable QTL that was identified by all three models using BLUPs (Figure 3) and nearly all individual environments was qDTA8, a large QTL on chromosome 8 that was strongly correlated with variation in days to anthesis as well as days to silking. The support interval for this QTL overlapped with two previously characterized flowering time QTL, *vgt1* and *vgt2*. As a well-studied, large effect QTL, *vgt1* provides a useful benchmark for comparison of the three allelic models.

At *vgt1*, the 16 founders are segregating (*MITE^+^/MITE^−^*) for the putative causal variant, a MITE insertion in a conserved non-coding sequence upstream of *ZmRap2.7* (Salvi *et al*. 2007; Castelletti *et al*. 2014). Four founders, B73, OH43, VA85, and B96, lack the MITE insertion, while the other 12 are *MITE^+^*. Looking at the most significant SNP for *qDTA8* from *GWAS_SNP_*, the alternate allele correlated imperfectly with the presence of the MITE in the founders (r=0.65). We expected to see *QTL_F_* effect sizes at this locus that match the allelic state of the founders, with *MITE^+^* founders having earlier effect sizes and *MITE^−^* founders having later effect sizes. However, for some founders the *QTL_F_* effect sizes at *vgt1* deviated from those expectations (Figure 4). Four *MITE^+^* founders, A632, F252, C103, and F492, had DTA BLUP effect size estimates later than the population average. While only F252 had a 95% confidence interval not overlapping zero, all had effect sizes significantly later than the other *MITE^+^* founders (t-ratio = 7.67, p-value <1e-4). This pattern was also seen in the effect sizes estimated in individual environments (Supplemental File 3). Lastly, at the most significant hit from *QTL_H_*, founders are grouped into haplotypes consistent with their allele at the MITE, but there are still far more than 2 distinct haplotypes (14). Analysis of the haplotype structure in the region around *vgt1* in the 16 founders showed clear differences between those that do and do not have the MITE insertion, but did not differentiate *MITE^+^* late founders from *MITE^+^* early founders (Figure S10).

**Figure 4.**
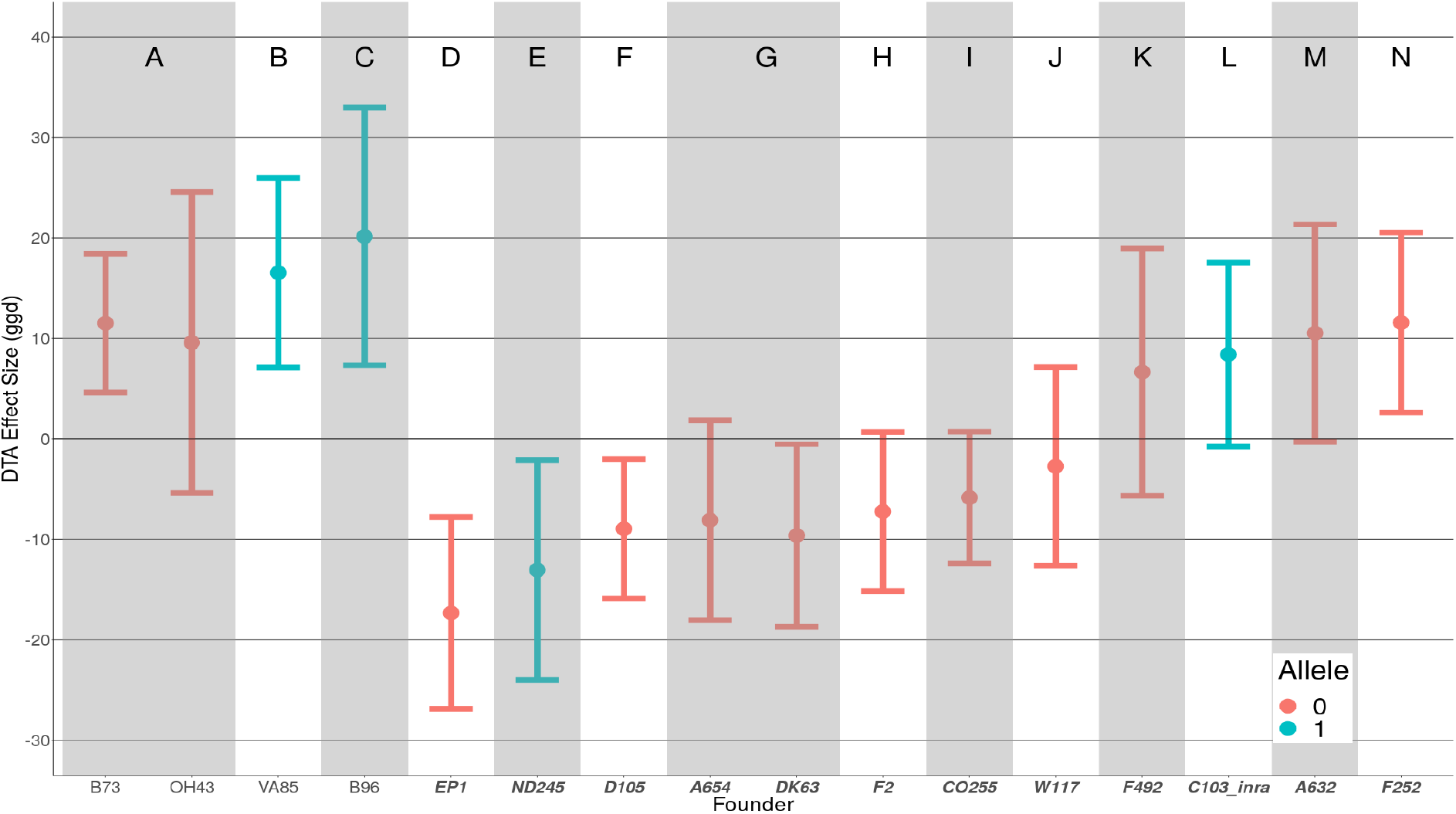
Estimated founder effect sizes for *vgt1*. Estimates of founder effect sizes relative to the population mean for Days to Anthesis (DTA) BLUP phenotypes using *QTL_F_*. Letters A-N and groupings of shaded blocks along the x-axis indicate founders that have shared haplotypes at the most significant *QTL_F_* SNP. Color indicates reference (0) and alternate (1) alleles at the most significant *QTL_F_* SNP. Positive effect sizes indicate later flowering and negative effect sizes indicate earlier flowering. Founders on the right (haplotypes D-N) labelled in bold and italics are lines that possess the MITE insertion (*MITE^+^*), while founders on the left (haplotyes A-C) lack the MITE insertion (*MITE^−^*).

One possible explanation for this observation is an epistatic interaction between *vgt1* and other loci in the genome. However, a genome scan for epistasis between *vgt1* and other loci did not yield any significant interactions (Figure S11). *QTL_F_* using only MAGIC lines predicted to have the MITE had two significant DTA BLUP QTL in the region of *vgt1* (Figure S12). One of these significant sites is located in close proximity to the causal gene for *vgt2, ZCN8*, and may be explained by this linked QTL (Guo *et al*. 2018). The second significant site is located 15Mb downstream of *vgt1*, suggesting that some local variation around the region of *vgt1* impacts the effect of the QTL on flowering time. This site may have an epistatic interaction with *vgt1* that did not pass the stringent genome-wide significance threshold. Alternatively, the relationship between the loci could be entirely additive, but the causal allele may only occur on the *MITE^+^* background.

### Founder Representation and Linkage Disequilibrium

Analysis of the MAGIC population showed that the overall representation of the 16 founders in the MAGIC DH lines was relatively even, with the highest percentage founder, A654, representing 6.7% and the lowest percentage founder, EP1, representing 5.2%, compared to the expectation of 6.25% for each founder (Figure 1C). However, multiple chromosome regions deviated significantly from the expected equal distribution (Figure 1B). Across individual regions of each chromosomes, 20.4% of the genome significantly deviated from null expectations compared to 100 simulated populations (5% significance threshold *χ*^2^ test p-value < 1.5e-09) (Figure S13). The fact that the *χ*^2^ test performed on 100 simulated MAGIC populations of 344 individuals resulted in far fewer significant *χ*^2^ peaks shows that the over- and under-representation of certain founder alleles was greater than would be expected by chance. It also shows that over- and under-representation of founder alleles in the population was not due to potential inaccuracy of R/qtl2 in assigning founders. These results suggests that a large amount of the over- and under-representation of founder alleles in the MAGIC population is biological, rather than a result of model error, and perhaps evidence of selection for or against particular founder alleles.

We estimated recombination rates and linkage disequilibrium across the MAGIC lines. The intra-chromosomal LD structure shows fast LD decay consistent with many recombination events (Figure S14). Unexpectedly, there was a large amount of high inter-chromosomal LD (Figure 5). Of a total of 9,796,630 SNP pairs with an *R*^2^ greater than or equal to 0.9, 365,345 (3.7%) of those pairs came from different chromosomes. The number of inter-chromosomal high LD regions was more than would be expected by chance: in 100 simulated populations, there were no SNP pairs with *R*^2^ greater than 0.9 detected between chromosomes. A large segment of inter-chromosomal LD between chromosome 3 and chromosome 8 (Figure 5) overlaps with the support intervals for *qDTA8* and *qDTA3-2*, corresponding to *vgt1* and *vgt3*, respectively, consistent with the possibility of selection driving differential representation of the founder alleles.

**Figure 5.**
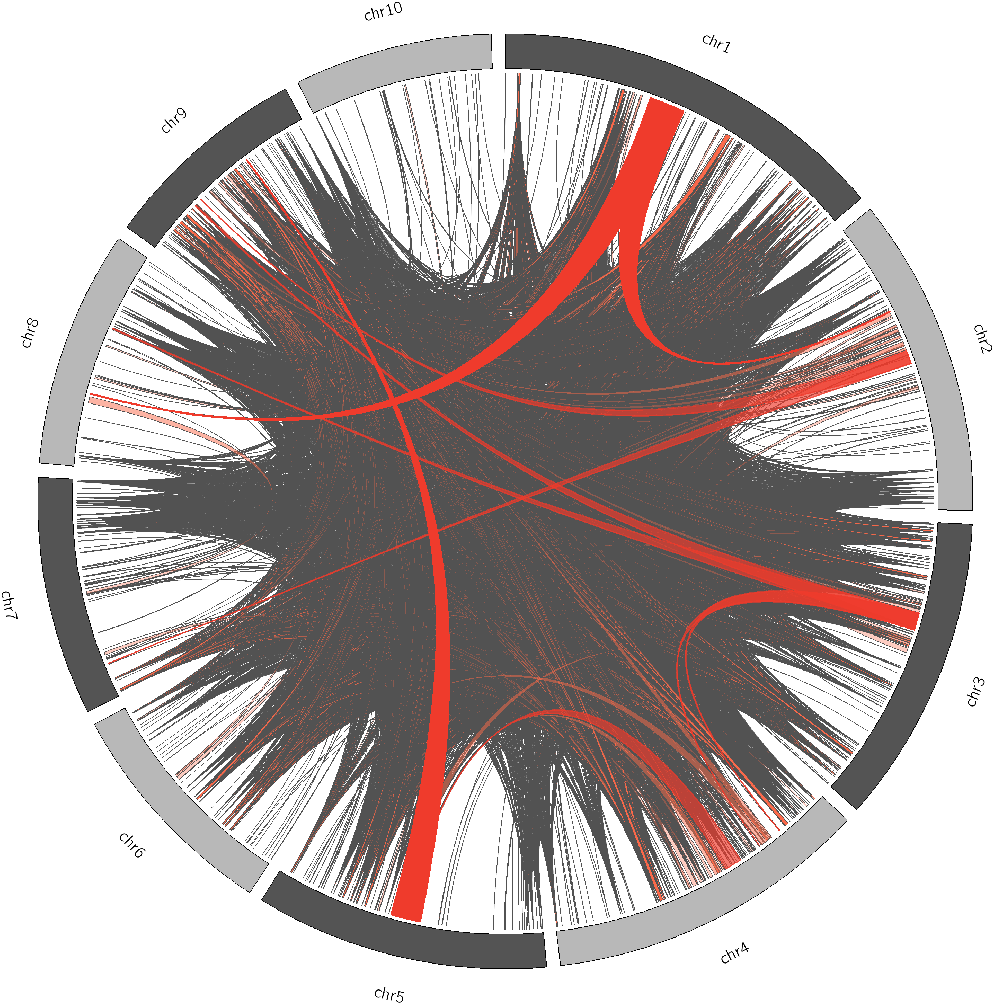
Inter-chromosomal Linkage Disequilibrium in the MAGIC Population. Ribbons represent regions of *R*^2^ > 0.8 between consecutive SNPs on different chromosomes. Dark, solid, red bands are regions larger than 20Mb on at least one of the chromosomes. Lighter, translucent red bands are regions greater than 1Mb and less than 20Mb. Grey bands are regions larger than 10kb.

Because individual DH lines were required to overlap in flowering time with the inbred tester in order to successfully make F1s for the MAGIC population, we hypothesized that the evidence of selection we saw in *χ*^2^ peaks and linkage disequilibrium might be due to selection on flowering time. In particular, the tester, MBS847, is a generally later flowering line and is *MITE^−^* at *vgt1* (Chardon *et al*. 2004), providing the opportunity for selection against early flowering alleles. As one test of this hypothesis, we asked whether genes involved in flowering were enriched in *χ*^2^ peaks, but found no evidence of enrichment (Figure S15).

## Discussion

We used three models of QTL allelic states to identify QTL in the MAGIC population, a bi-allelic model (*GWAS_SNP_*), a founder multi-allelic model (*QTL_F_*), and an ancestral haplotype mult-allelic model (*QTL_H_*). The *GWAS_SNP_* method should be most powerful at identifying QTL for which the causal variant is bi-allelic and the tagged SNP is in tight LD with the causal variant. However, for multi-allelic QTL or QTL for which LD is low between tagged SNPs, this method should have lower power. *QTL_F_*, which assumes that all founders possess distinct alleles, increases the odds of detecting both QTL that are multi-allelic and QTL whose causal variant is not in tight LD with any one tagged SNP. And while the higher number of parameters that must be fit by this model may also reduce power because the regions tested are much larger than one SNP, it also reduces the multiple testing burden. Lastly, *QTL_H_* potentially improves on the power of *QTL_F_* to detect QTL that meet the above criteria by reducing the number of parameters that must be estimated. There is, however, the potential for *QTL_H_* to obscure the signal of some QTL if founders are called as having the same ancestral haplotype when they actually differ for a causal variant. Due to the fact that *QTL_H_* and *QTL_F_* only take into account recent recombination events, whereas *GWAS_SNP_* uses historical recombination, we predicted that *GWAS_SNP_* would result in higher resolution of QTL support intervals. Higher resolution QTL are ideal in that they make it easier to narrow down candidate genes and potential causal variants.

The results of using these three models of genetic architecture to identify QTL suggest that each has its own advantages and disadvantages in terms of how many and which QTL they can identify. Overall, the *QTL_F_* model performed the best in terms of the number of QTL identified, although there were multiple QTL identified uniquely in all models. The larger size of *GWAS_SNP_* support intervals was unexpected, as this method is often used to fine map QTL regions identified by linkage mapping. We suspect that this finding is either the result of the somewhat naive method used to determine QTL support intervals, or residual long-distance LD caused by the funnel crossing scheme. A twofold drop in log_10_ p-values heavily penalizes QTL that just pass significance, and the 100Mb window is blind to QTL peaks that contain multiple QTL. As a result, it is difficult to determine a reason for the difference in support interval size between the models.

Previous studies have used variations of these methods to identify QTL, and some have directly compared them. The use of combined linkage and association analyses, sometimes referred to as linkage disequilibrium - linkage association (LDLA) was first proposed by Meuwissen and Goddard (2001), who used predicted IBD probabilities between parents using an evolutionary model and applied them to linkage mapping. LDLA has been used in multiple studies to enhance QTL detection in multi-parent populations in maize (Giraud *et al*. 2017; Yu *et al*. 2008; McMullen *et al*. 2009) and other organisms (Hérault *et al*. 2018). Jansen *et al*. (2003) used a haplotype-based method for QTL mapping and showed through simulation that this strategy could reduce the number of estimated parameters and, therefore, increase power. Different means of determining ancestral haplotype blocks from parental sequences have been used, with clusthaplo (Leroux *et al*. 2014), an extension of the software MC-QTL (Jourjon *et al*. 2004), being a commonly used algorithm in recent studies. Bayesian frameworks have also been implemented in real (Pérez-Enciso 2003) and simulated (Bink *et al*. 2012) multi-parent populations. Giraud *et al*. (2014) used both a haplotype- and founder-based approach in two nested association mapping populations of Northern European flint and dent maize lines created by Bauer *et al*. (2013) and genotyped with a 50K SNP array (Ganal *et al*. 2011). Giraud *et al*. (2014) used clusthaplo to determine haplotype blocks based on IBD between parents and used discrete founder and haplotype values in their models.

Previous studies have compared similar models differing in a few notable ways. The type of multi-parent population used in studies has varied across studies. To our knowledge, comparison of the performance of these models in a MAGIC population has not been done. In addition, our method of generating haplotype blocks in *QTL_H_* is distinct from previous work. Lastly, the use of the package GridLMM (Runcie and Crawford 2019) allowed us to use continuous rather than discrete representations of founder and haplotype state, which allowed for the incorporation of uncertainty due to genotyping or model error into our tests for association.

Interestingly, the performance of bi-allelic, founder, and ancestral haplotype models differs across studies. Giraud *et al*. (2014) found that their haplotype model outperformed the founder and SNP models in terms of the number of QTL identified using EU-NAM Flint and Dent maize populations. In contrast, Garin *et al*. (2020) found that in the EU-NAM Flint population, the bi-allelic model detected a larger number of unique QTL, compared to parental or ancestral haplotype models. Bardol *et al*. (2013) found that in two multi-parent dent populations, their bi-allelic model and ancestral haplotype model generally outperformed the parental linkage model, although benefits of these models varied by dataset. The performance of the three models seems to depend heavily on the diversity of the parents used to generate the population. For populations with more diverse founders, it would be expected that there would be fewer shared haplotypes between founders, reducing the efficacy of a haplotype model (Giraud *et al*. 2014). The fact that the *QTL_F_* model outperformed the *QTL_H_* and *GWAS_SNP_* in our population suggests that the MAGIC population contains a relatively more diverse representation of European and North American flint and dent than populations used in previous studies. It is also possible that the structure of multi-parent populations has an effect on the performance of the three models, compared to previous studies which used nested association mapping (Giraud *et al*. 2014; Garin *et al*. 2020) and factorial populations Bardol *et al*. (2013).

Differences in the estimated effect sizes across models offer suggestions as to the reason for their differences in QTL detection. QTL that were only found in the *GWAS_SNP_* method most likely have a bi-allelic causal variant. It is likely that the increased number of parameters in the *QTL_F_* and *QTL_H_* models reduce statistical power when the true number of functional alleles is low. Similarly, we predict multi-allelic QTL were more likely to be identified by the *QTL_F_* or *QTL_H_* models and not the *GWAS_SNP_* method unless the effect size was large. For QTL that were identified in the *QTL_H_* method and not the *QTL_F_* method, there tended to be a lower number of unique ancestral haplotypes, suggesting that *QTL_H_* was more successful in finding these QTL due to improved power when there were fewer functional alleles than founders. For QTL that were not identified by the *QTL_H_* method, particularly for QTL that were successfully identified by *QTL_F_*, there are two possible explanations. Because there were more regions tested for *QTL_H_* than for *QTL_F_*, if the true number of ancestral haplotypes at the QTL is large, the *QTL_H_* method may actually have lower power than *QTL_F_* because the number of tests is higher. Alternatively, it may be the result of a failure of *QTL_H_* to accurately represent the true haplotype structure of the QTL region.

On the whole, most QTL were found by all three methods, so there was limited ability to draw reliable conclusions about underlying mechanisms that caused the methods to perform better or worse. Generally, the comparison of QTL detection and effect size estimates suggested that the methods failed and succeeded on a QTL-by-QTL basis. This is to be expected, as each QTL is the result of a distinct set of one or more causal alleles with a unique evolutionary history and pattern of linkage disequilibrium within the population.

Whether QTL that appeared in only one method are due to false positives or true differences in the methods’ abilities to identify QTL with different genetic architectures cannot be determined, but many of the QTL identified in the MAGIC have underlying candidate genes or have been found in previous studies, providing support to their biological reality. Multiple flowering time QTL support intervals overlap or are close by previously identified flowering time genes and QTL. *qDTA9* is nearby the previously identified maize flowering time gene *ZmCCT9* (Huang *et al*. 2018). *qDTA3-2* overlaps with *vgt3*, whose underlying gene was identified as *ZmMADS69* (Liang *et al*. 2019). *qDTA3-1* is nearby a recently identified flowering time QTL also associated with phasphatidylcholine levels (Rodríguez-Zapata *et al*. 2021). The support interval for *qDTA8* overlaps with two flowering time QTL, *vgt1*, which we discuss in length, and another, *vgt2*. The causal gene for *vgt2* is *ZCN8*, which is the maize ortholog of FT in *Arabidopsis* (Lazakis *et al*. 2011). Variation in the promoter region of *ZCN8* between temperate maize and teosinte suggests that earlier flowering alleles were under selection during the process of maize domestication (Guo *et al*. 2018; Bouchet *et al*. 2013). It is interesting to note that there is strong overlap in the support intervals of QTL found on chromosome 3 between flowering time and harvest grain moisture (Figure 3), perhaps due to developmental pleiotropy linking flowering time and the moisture of kernels at harvest. Three GxE QTL were detected in this population (Hudson *et al*. in prep) using the *QTL_F_* model, but none overlapped with the main effect QTL we detected in this study. One of these QTL appeared to be a false positive resulting from low representation of one of the founders in the region, indicating the potential for low founder sample size to confound QTL results.

Due to the fact that the MAGIC population is segregating for *vgt1*, it provides an opportunity to further study the mechanism behind the QTL’s affect on flowering time. One benefit of using founder and haplotype approaches lies in the potential to dissect the effects of individual founders and haplotypes within QTL. This allowed us to look more closely at *vgt1* and observe an interesting pattern of effect sizes that deviated from our expectations based on previous research. Previous research has shown that variation in flowering time at this site is strongly correlated with a MITE insertion about 70kb upstream of the flowering time regulator, *ZmRAP2.7*, an APETALA-like transcription factor, with the presence of the MITE associated with an earlier flowering time (Castelletti *et al*. 2014). Within maize heterotic groups, Flint maize lines tend to possess the early-flowering allele of *vgt1* (*MITE+*), while dents (such as B73) tend to carry the late-flowering allele (*MITE-*) (Salvi *et al*. 2007). In addition to being a crucial agronomic trait, flowering time contributes to local adaptation for annual plants such as maize, ensuring that individuals can reproduce within the growing season of their environments. The frequency of the MITE in maize populations follows a latitudinal gradient, suggesting that the early *MITE^+^* allele was selected for during the process of maize adaptation to temperate climates (Navarro *et al*. 2017). It has also been shown that there are differentially-methylated regions around *vgt1* between B73, landrace maize, and teosinte (Xu *et al*. 2020). The hypothesized mechanism of action is that the MITE represses expression of the negative flowering time regulator, *ZmRAP2.7*, possibly due to changes in methylation around the insertion, resulting in earlier induction of flowering (Castelletti *et al*. 2014). However, the MITE has not yet been experimentally shown to result in earlier flowering. A recent study using multiple multiparent populations suggested that variation in the effect of *vgt1* in different genetic backgrounds was due to local genetic variation surrounding *vgt1*, rather than epistasis with distant loci (Rio *et al*. 2020). The observed lack of significant epistasis with *vgt1* in this study (Figure S13), combined with our results of *MITE^+^ QTL_F_* (Figure S11), appear consistent with this idea. This finding suggests two possibilities: either (1) that the causal variant underlying *vgt1* is some as-yet unidentified variant that is in tight, but imperfect linkage disequilibrium with the MITE insertion, or (2) that the MITE insertion directly impacts flowering time, and that another variant nearby has a modifying effect on the MITE. This opens up new areas of inquiry for future studies.

The population displayed unexpected patterns of uneven founder representation along the genome (Figure 1C). The apparent high over- or under-representation of some founder alleles may be due to these founders being in close IBD with another founder, resulting in uncertainty in the founder probabilities. Nonetheless, the high accuracy of founder assignment and low number of sites that significantly deviate from equal founder distribution in our simulations make this explanation seem unlikely. If we assume that the simulated populations are accurate representations of the construction of the MAGIC population without selection, this result would suggest that the differences in founder representation observed in the actual population may be due to selection for or against certain founder alleles. The complex crossing scheme has the potential to introduce the influence of selection, which may reduce the genetic variation that we are attempting to study.

Another interesting observation of the population was the relatively high levels of inter-chromosomal LD (Figure 5), which deviated significantly from that obtained from simulations. A potential consequence of inter-chromosomal LD is the chance for confounding of association analyses, namely resulting in the detection of “ghost” QTL. Although for the discussed QTL detected that were in high inter-chromosomal LD with one another, *qDTA8* and *qDTA3-2*, their effects on flowering time were independent, the chance for false positives and inaccurate support intervals due to LD structure is still worth noting. Inter-chromosomal LD has been detected in multiple populations of domesticated organisms, where breeding has resulted in the preservation of certain combinations of favorable alleles between chromosomes (Robbins *et al*. 2010; Malysheva-Otto *et al*. 2006). Strong selection and positive or negative epistasis in natural populations have also been shown to create a pattern of inter-chromosomal LD (Kulminski 2011; Gupta *et al*. 2021; Hench *et al*. 2019; Petkov *et al*. 2005). Both of these observations suggest that forces other than those of random segregation have operated on the MAGIC population.

## Conclusion

The MAGIC population presented here provides a useful resource for investigating quantitative trait variation in temperate maize. As a multi-parent population, it has the advantages of increased diversity compared to bi-parental mapping populations, no population structure, and higher power to detect QTL with lower allele frequencies. Simulations of the MAGIC population provide an opportunity to validate assignment of founder identities, as well as generate null expectations for various aspects of the population. Overall, we find that a founder multi-allelic model identifies the most QTL, although all three models of allelic state are effective at identifying QTL. The benefits of increasing statistical power by reducing model parameters in bi-allelic and ancestral haplotype models seem to be tempered by the true allelic complexity of the multi-parent population being studied. We conclude that if the goal of a study is to find as many QTL as possible, then it would be most useful consider the genetic diversity contained within the population in choosing a model in order to maximize QTL identification.

## Acknowledgments

S.G.O. was supported by the UC Davis Dept. of Plant Sciences and NSF grant 1754098. A.I.H. was supported by National Science Foundation Graduate Research Fellowship 1650042. J.R.I. was supported by USDA Hatch project CA-D-PLS-2066-H 548. D.E.R. was supported by USDA Hatch project 1010469 and USDA NIFA grant no. 2020-67013-30904. We would like to acknowledge the Limagrain breeding support and trialing team for producing the doubled haploids and running the field trials used in this study.

## Supplement

**Figure S1.**
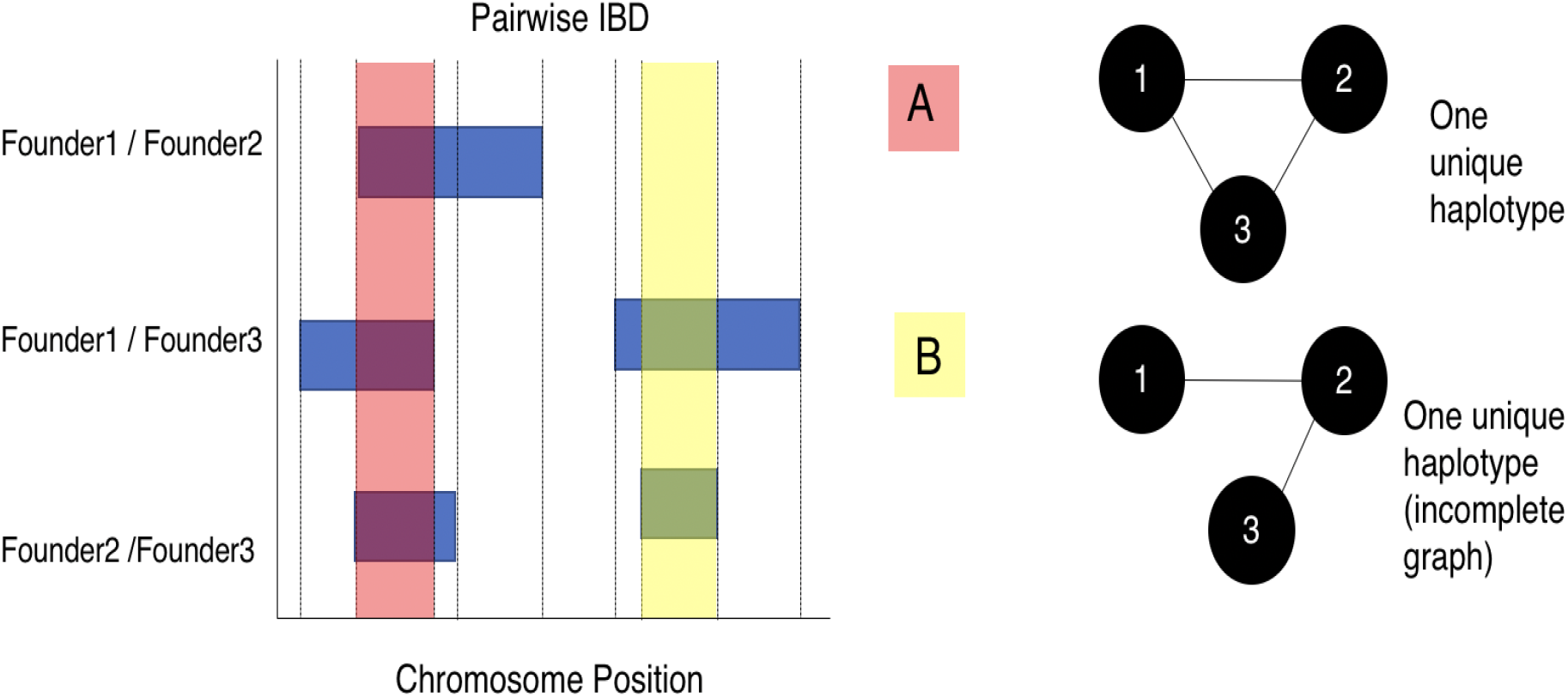
Complete and Incomplete IBD Graphs. **(left)** An example of the creation of haplotype blocks. The presence of pairwise IBD between two founders along the chromosome is shown as blue blocks. Black verticals show the demarcation of distinct haplotype blocks at the start or and of a new pairwise IBD segment. The red and yellow shaded areas show examples of complete and incomplete founder IBD graphs, respectively. **(right)** A complete **(A)** and an incomplete **(B)** founder IBD graph. In cases where haplotypes were incomplete, the missing edges (i.e. the edge between Founder 1 and Founder 3) were filled in for downstream analysis.

**Table S1.** Size of QTL Support Intervals By Model.

| Model | Avg. Size (Mb) | Median (Mb) | Size | Std. Dev (Mb) | Avg. Size (cM) | Median (cM) | Size | Std. Dev (cM) |
| --- | --- | --- | --- | --- | --- | --- | --- | --- |
| $GWAS_{SNP}$ | 17.53 | 19.67 | | 10.19 | 7.31 | 8.21 | | 3.50 |
| $QTL_F$ | 14.99 | 10.28 | | 13.64 | 6.22 | 5.88 | | 3.78 |
| $QTL_H$ | 13.72 | 9.24 | | 10.27 | 5.75 | 5.45 | | 3.63 |

**Figure S2.**
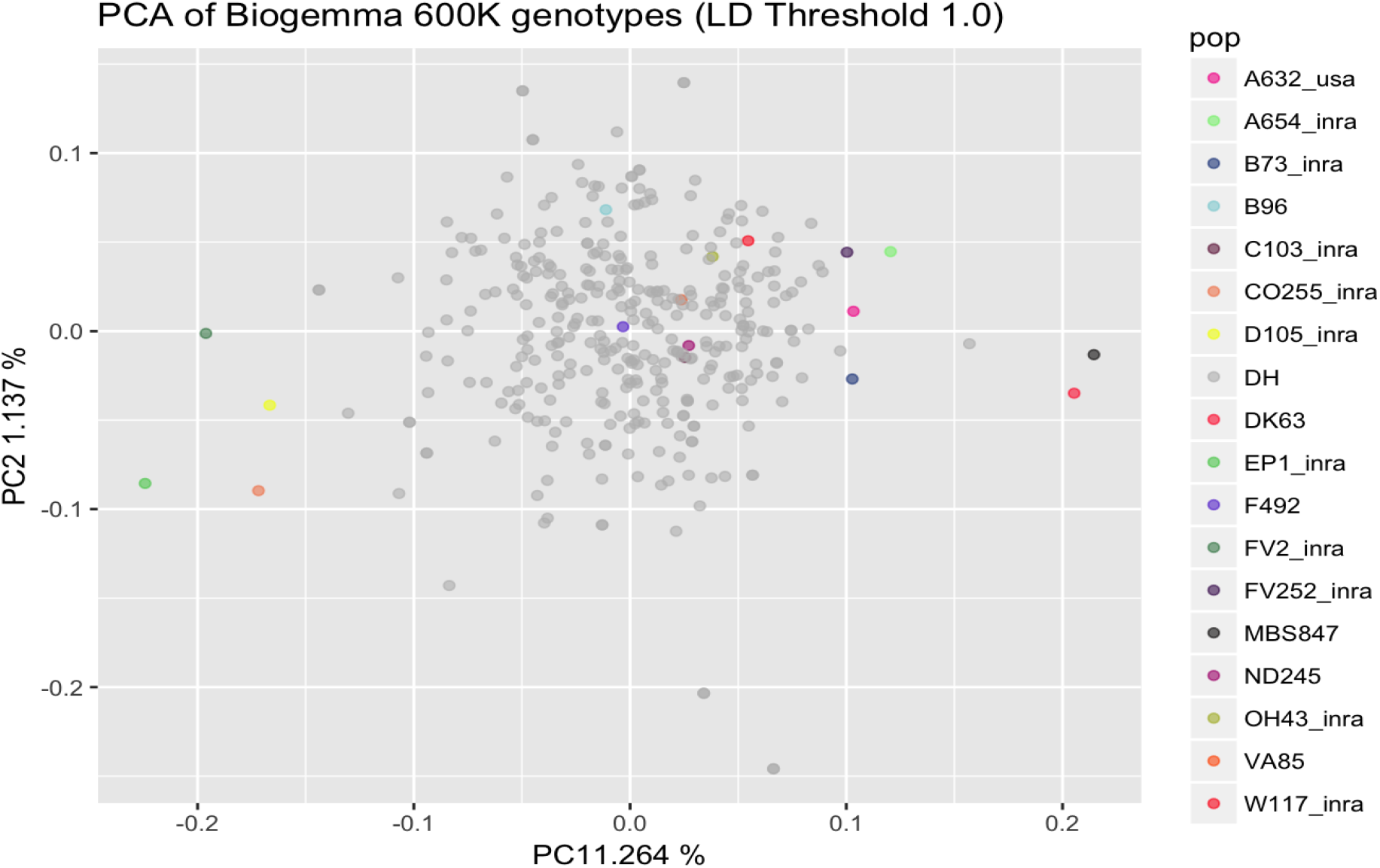
Principal Component Analysis of the 16 Founders, Tester, and the 344 MAGIC DH Lines. 600K genotype data was filtered using an LD filter of 1 cM. The points in grey are MAGIC lines. The colored points show the tester, MBS847 (dark grey) and the 16 founders.

**Figure S3.**
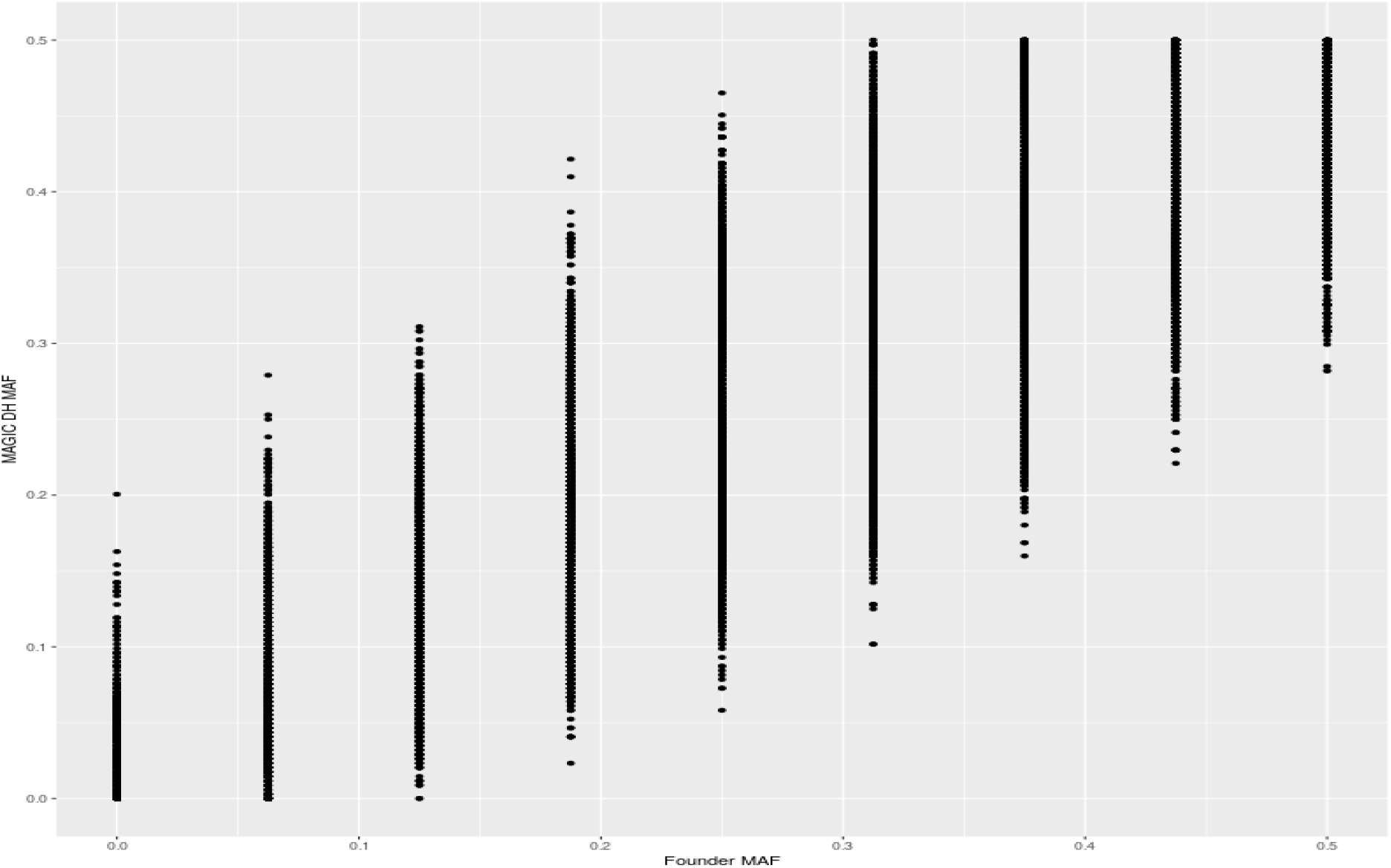
Minor Allele Frequencies of SNPs in the 16 Founders and the MAGIC lines. The x-axis shows the minor allele frequency of 600K SNPs in the 16 founders. The y-axis shows the minor allele frequency of the corresponding SNPs in the MAGIC population.

**Figure S4.**
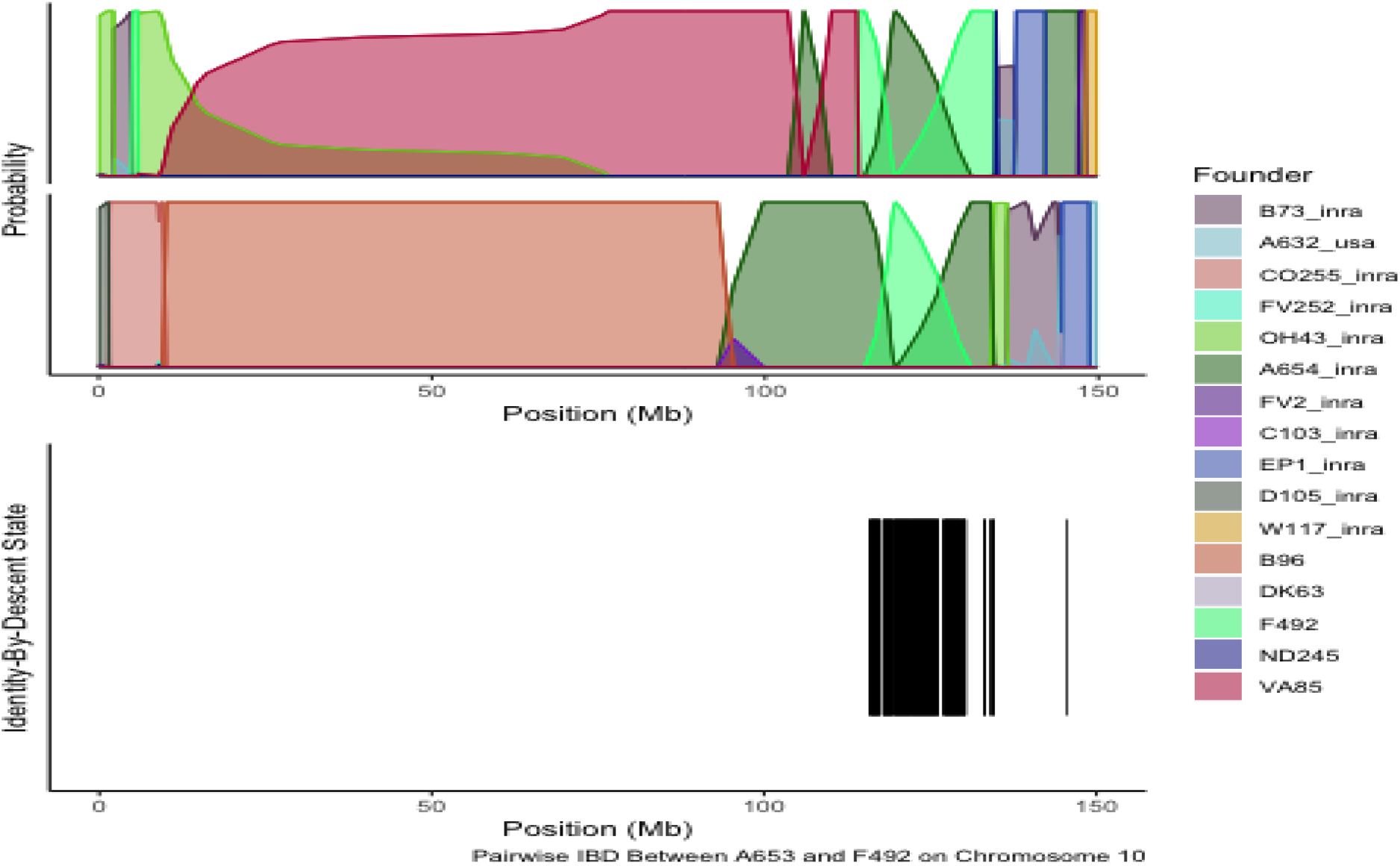
Overlap between uncertainty in founder probabilities and pairwise Identity-By-Descent between founders. **(top)** An example of founder probabilities across chromosome 10 for two MAGIC lines. The y-axis shows the probability of the line being derived from each of the 16 founders, shown in different colors. **(bottom)** Pairwise IBD state across chromosome 10 between two founders, F492 and A654, which correspond to uncertainty in founder probabilities above.

**Figure S5.**
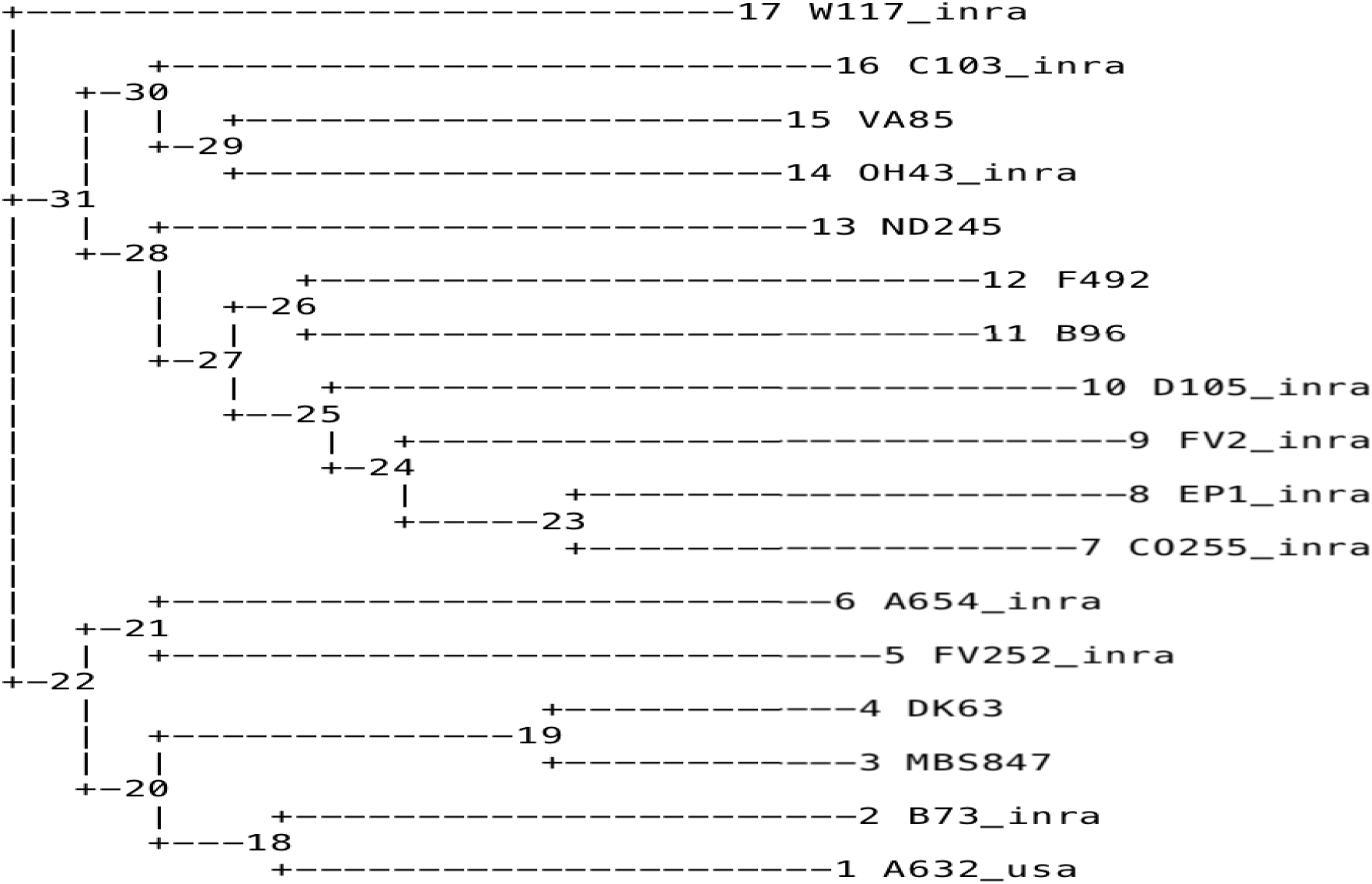
Neighbor-Joining Tree of the 16 Founders and Tester using the 600K genotype data. A UPGMA dendrogram created using the TASSEL software (Bradbury *et al*. 2007).

**Figure S6.**
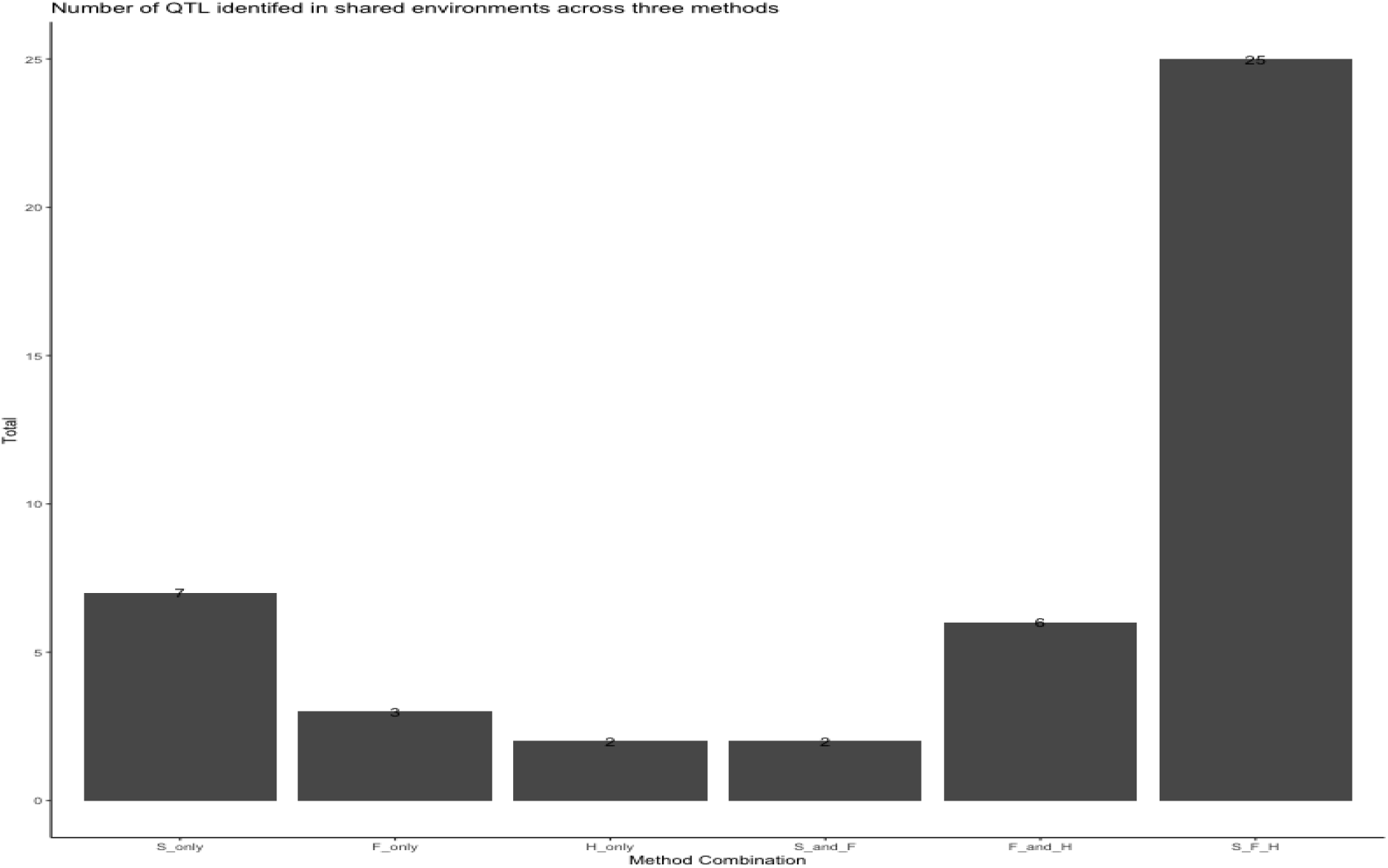
Number of Environment-Specific QTL Found by Method. S: *GWAS_SNP_*, F: *QTL_F_*, H: *QTL_H_*. Environment-specific QTL are QTL for a phenotype that were identified in one environment. QTL were called as found by multiple methods if their support intervals overlapped (see Methods).

**Figure S7.**
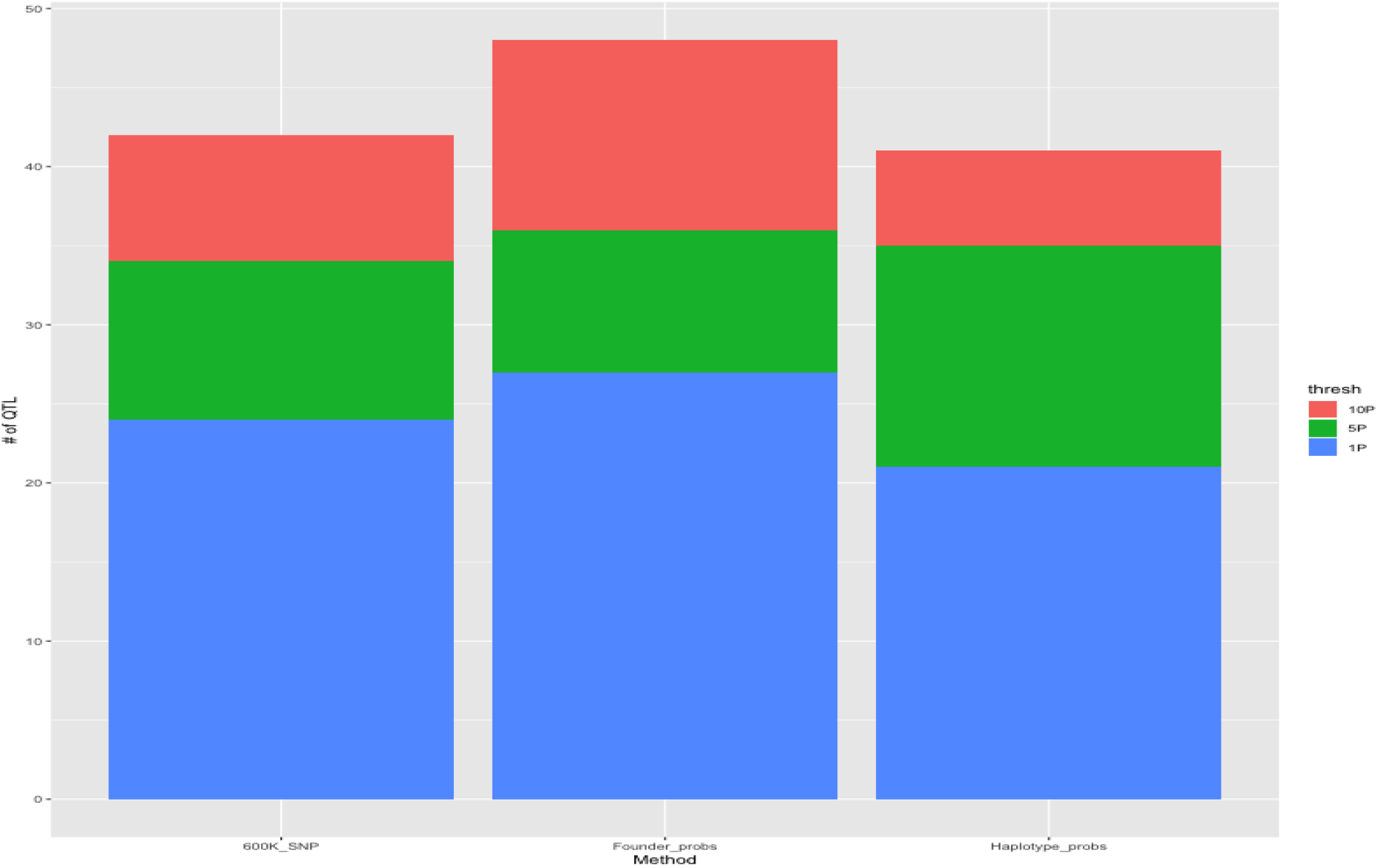
Total Number of Environment-QTL Found By Method By Significance Threshold. S: *GWAS_SNP_*, F: *QTL_F_*, H: *QTL_H_*. Environment-specific QTL are QTL for a phenotype that were identified in one environment. Color indicates new QTL found by raising the significance threshold from 1% (red) to 5% (green) to 10% (blue).

**Figure S8.**
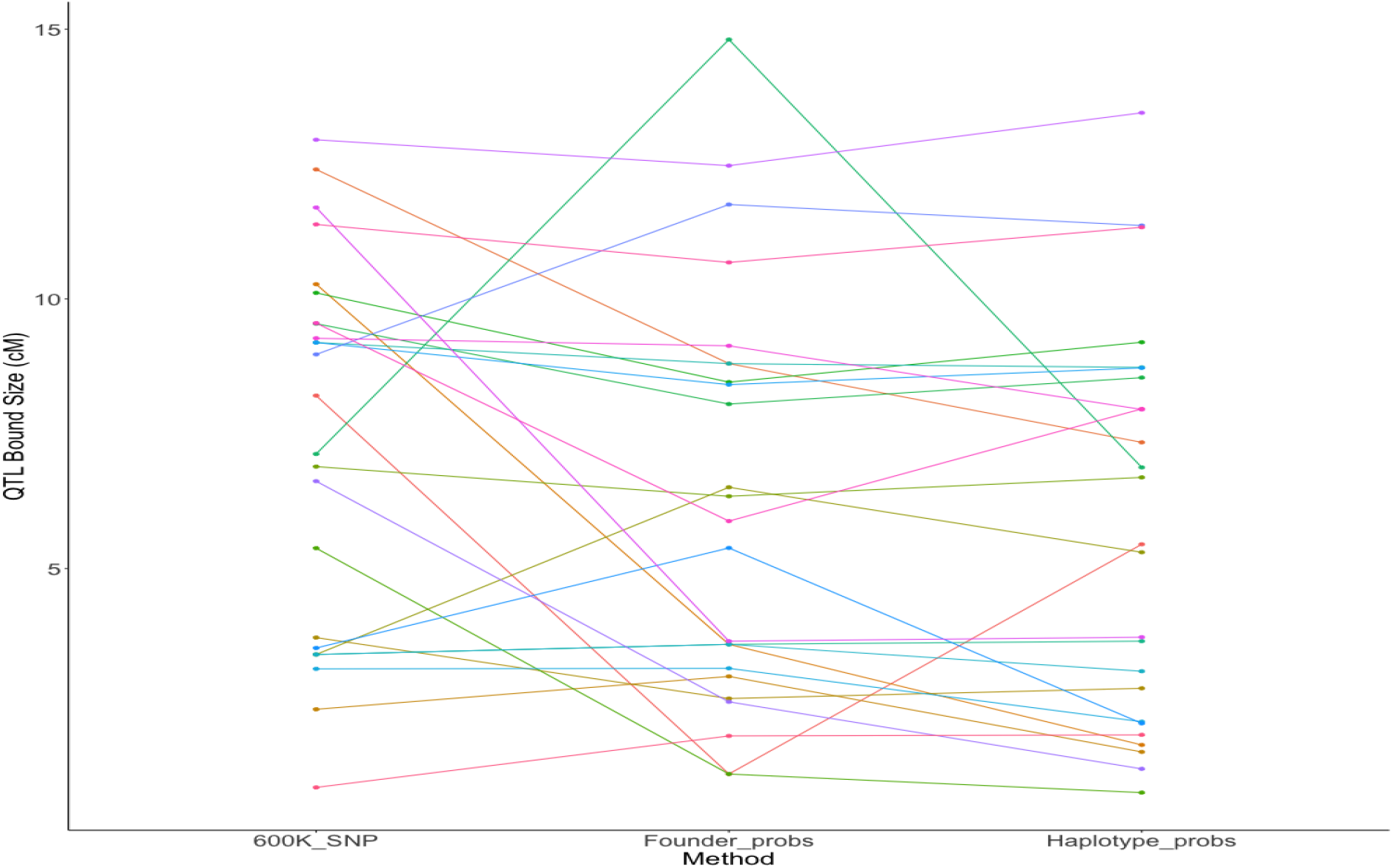
Size of QTL Support Intervals for Shared QTL by Method in Genetic Distance (cM) 600K_SNP: *GWAS_SNP_*, Founder_probs: *QTL_F_*, Haplotype_probs: *QTL_H_*. Colors indicate individual Environment-QTL.

**Figure S9.**
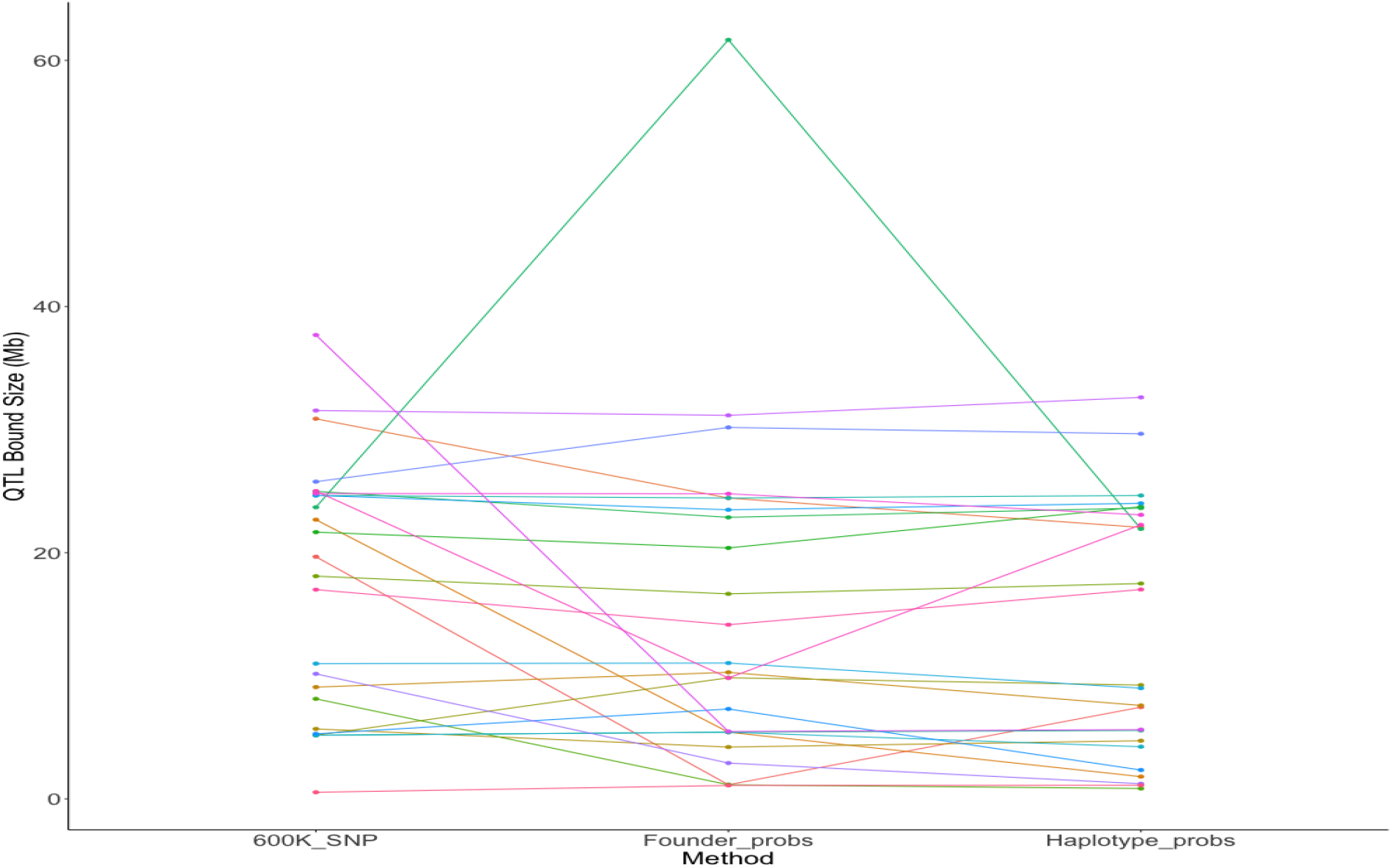
Size of QTL Support Intervals for Shared QTL by Method in Physical Distance (Mb) 600K_SNP: *GWAS_SNP_*, Founder_probs: *QTL_F_*, Haplotype_probs: *QTL_H_*. Colors indicate individual Environment-QTL.

**Figure S10.**
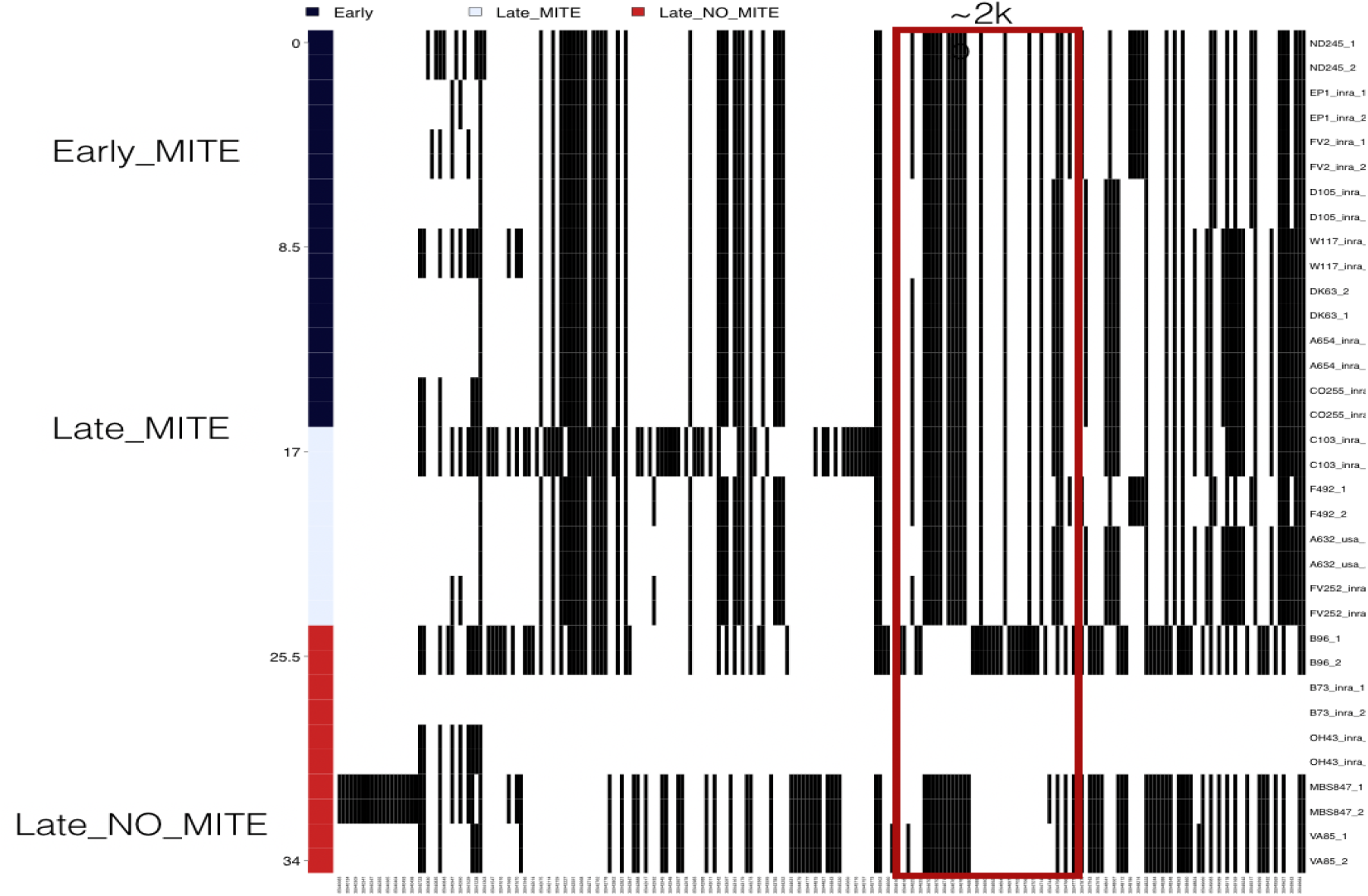
Haplotype View of the MITE insertion underlying *vgt1* in the 16 Founders and tester, MBS847. Red blocks indicates the location of the MITE insertion. Founders are separated by *MITE^-^*, *MITE^+^* Early, and *MITE^+^* Late based off of their allele at *vgt1* and groupings based on the estimated founder effect size. Created using the haplostrips software (Marnetto and Huerta-Sánchez 2017)

**Figure S11.**
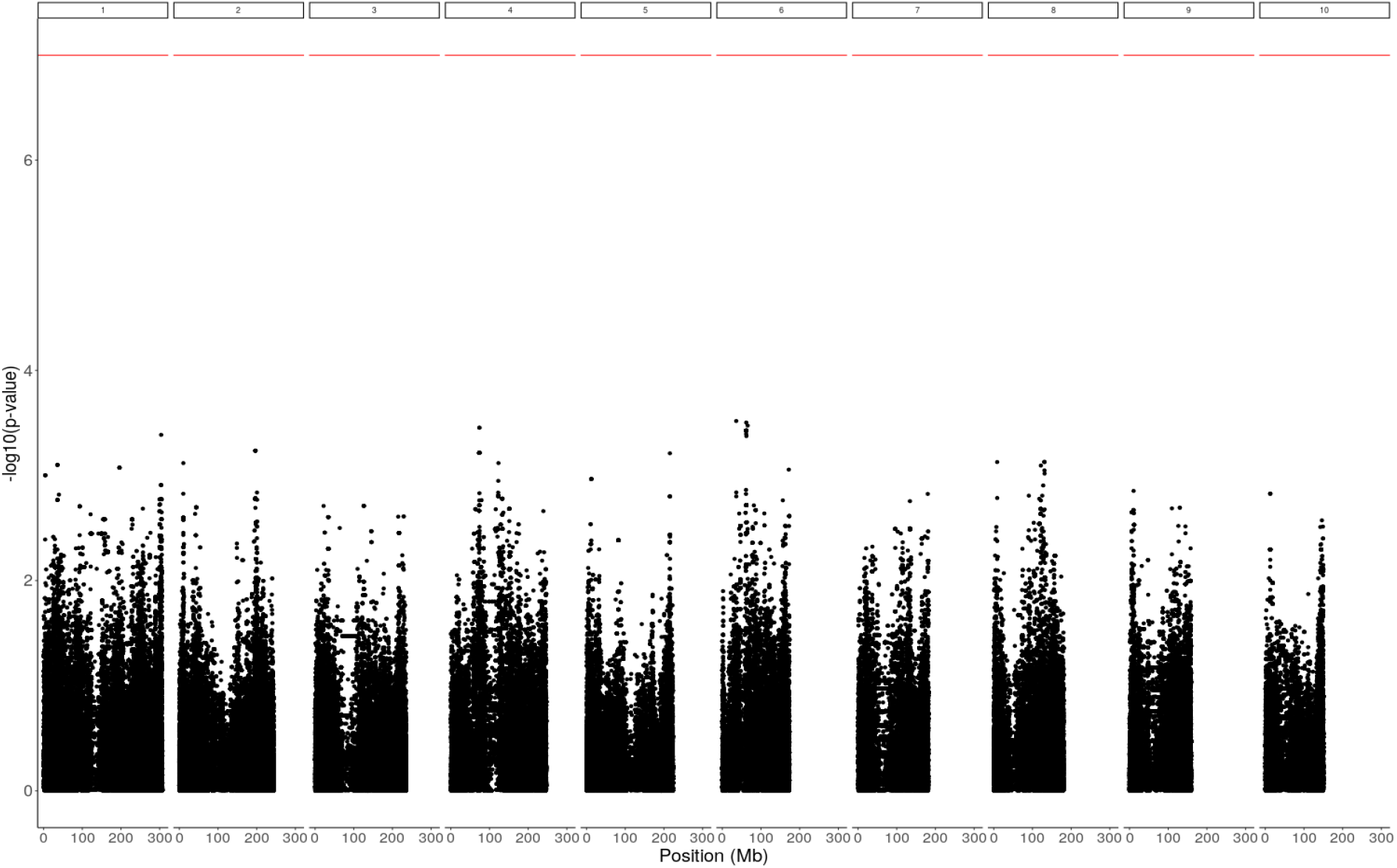
Genome Scan for Epistasis with *vgt1*. Red line indicates a Bonferroni 5% significance threshold.

**Figure S12.**
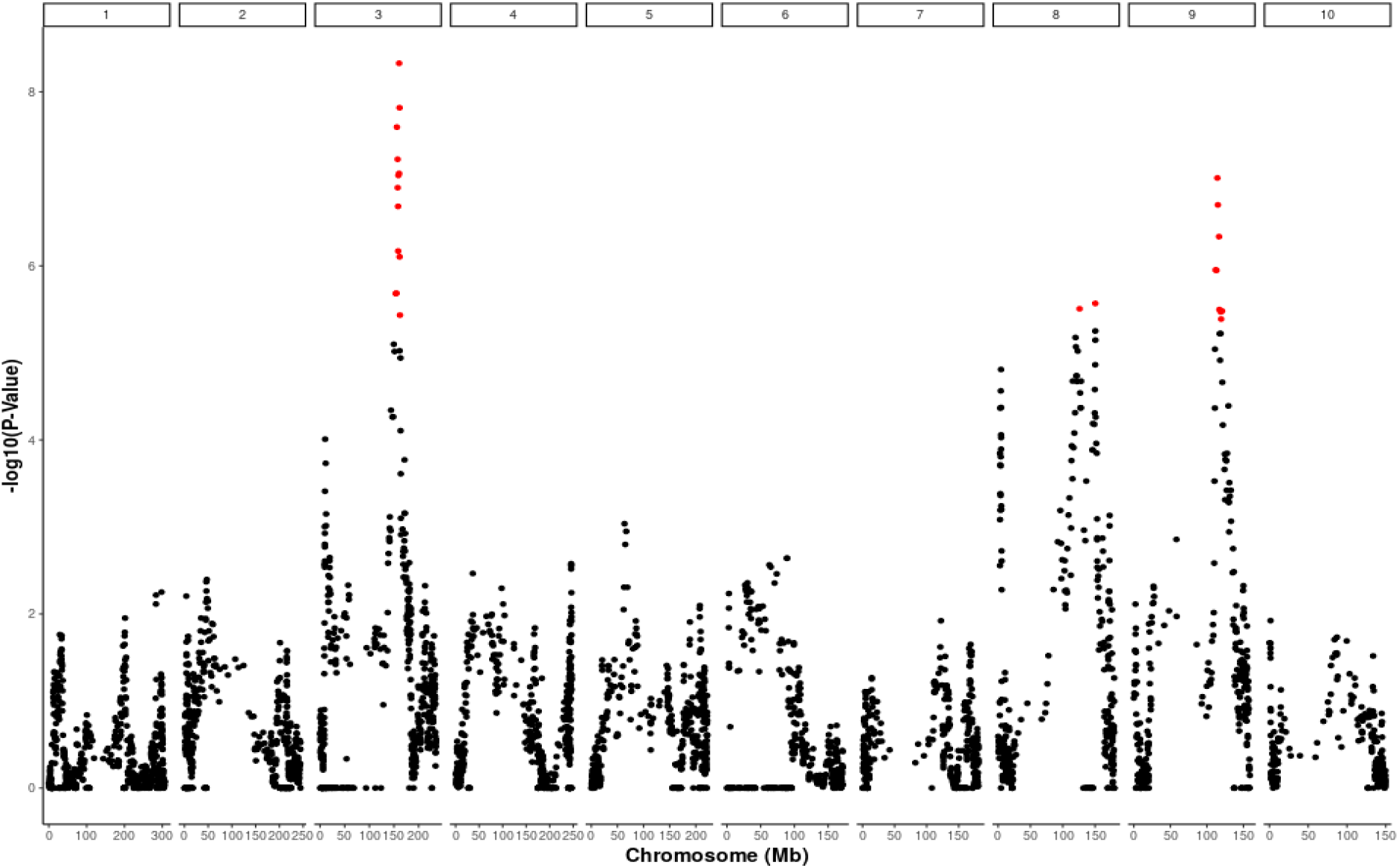
Manhattan plot Days to Anthesis BLUPs using only *MITE^+^* MAGIC lines and *QTL_F_*. Red points indicate SNPs above a 5% permutation significance threshold.

**Figure S13.**
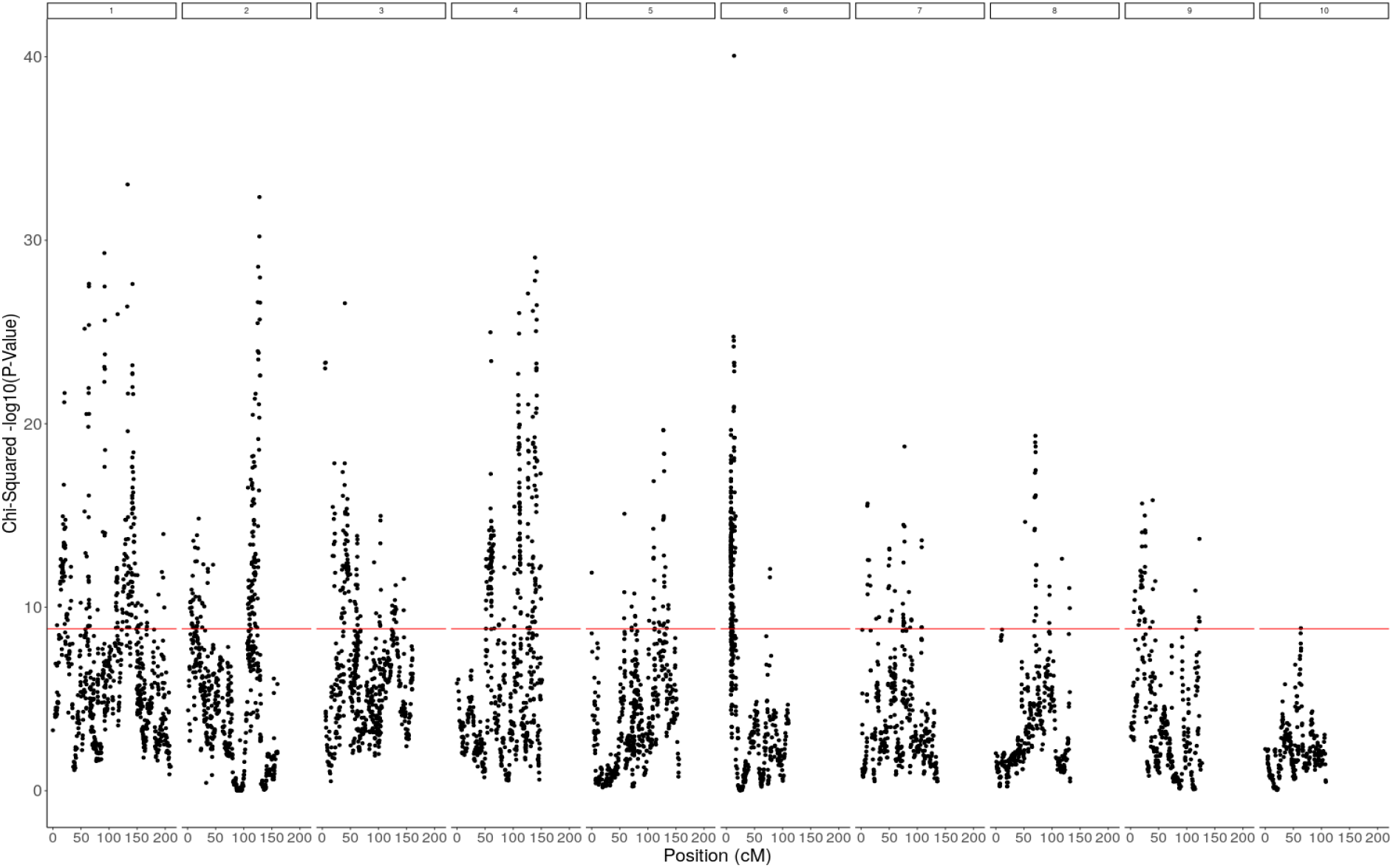
*χ*^2^ Peaks for Deviation from Equal Founder Distribution by Chromosome. Red line indicates a 5%significance threshold (-log10(p-value=8.82)) obtained from 100 simulated populations.

**Figure S14.**
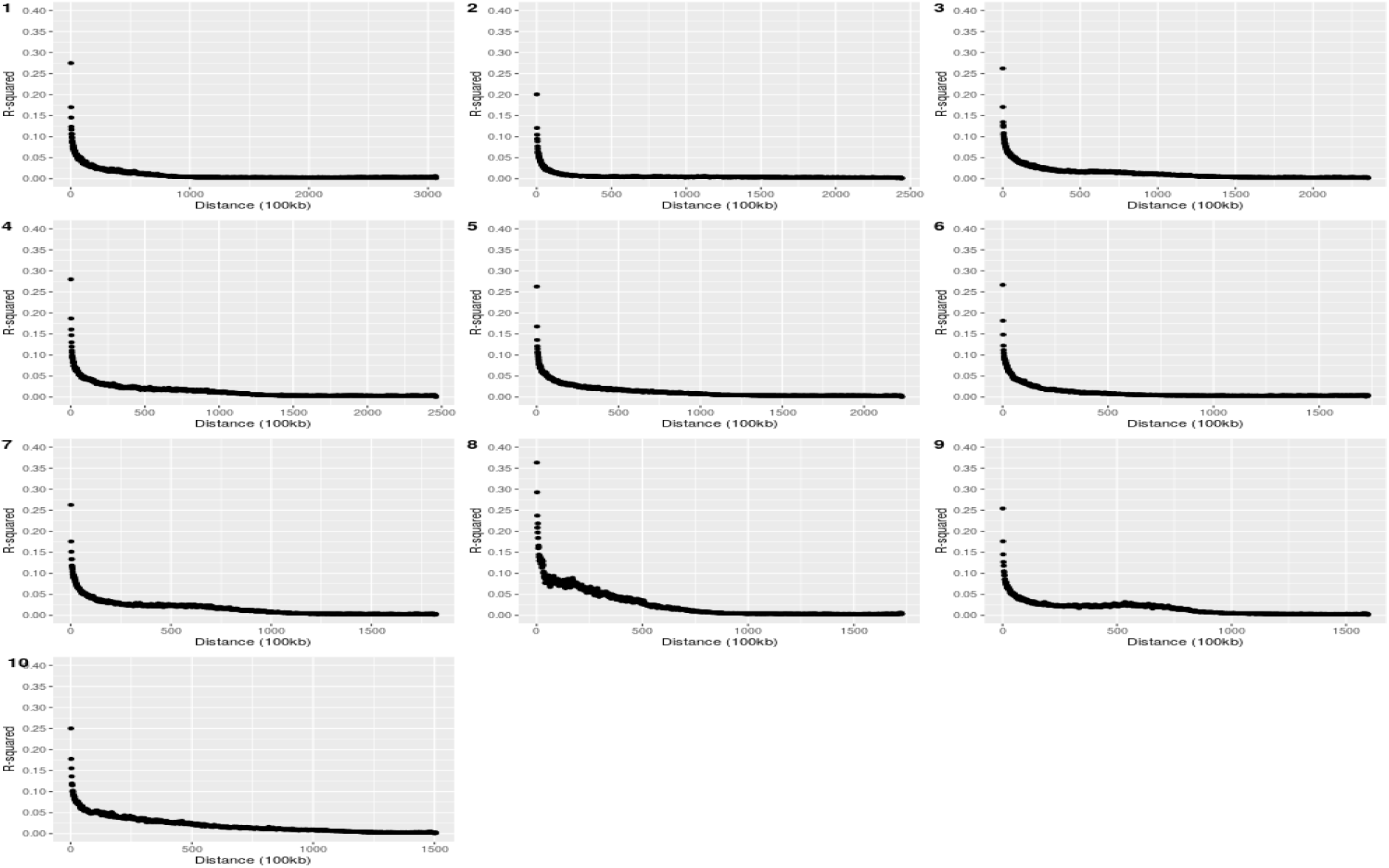
Linkage Disequilibrium of the MAGIC population by Chromosome. Average *R*^2^ of pairwise SNPs broken into bins of 100kb by distance.

**Figure S15.**
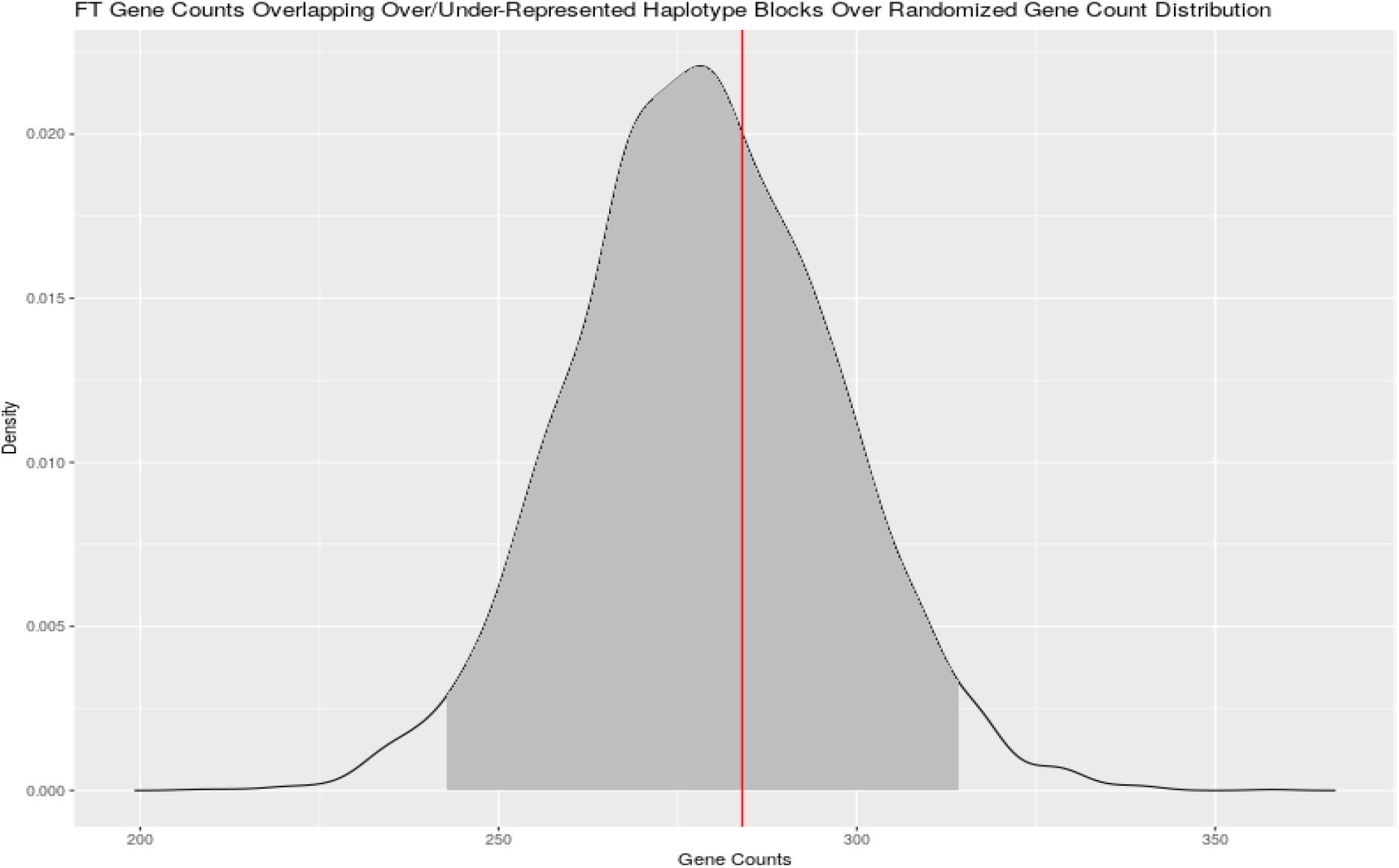
Enrichment of Flowering Time Genes in *χ*^2^ Peaks. Density plot of 1,000 permutations of number of 904 randomly selected genes that overlap with *χ*^2^ peaks for over- or under-representation of haplotypes in the MAGIC population. Red line indicates the actual number (284) of FT genes from the list of 904 that overlapped with *χ*^2^ peaks.

